# Multi-Omics after O-GlcNAc Alteration Identifies Cellular Processes Working Synergistically to Promote Aneuploidy

**DOI:** 10.1101/2024.04.16.589379

**Authors:** Samuel S. Boyd, Dakota R. Robarts, Khue Nguyen, Maite Villar, Ibtihal Alghusen, Manasi Kotulkar, Aspin Denson, Halyna Fedosyuk, Stephen A. Whelan, Norman C.Y. Lee, John Hanover, Wagner B. Dias, Ee Phie Tan, Steven R. McGreal, Antonio Artigues, Russell H. Swerdlow, Jeffrey A. Thompson, Udayan Apte, Chad Slawson

## Abstract

Pharmacologic or genetic manipulation of O-GlcNAcylation, an intracellular, single sugar post-translational modification, are difficult to interpret due to the pleotropic nature of O-GlcNAc and the vast signaling pathways it regulates. To address this issue, we employed either OGT (O-GlcNAc transferase), OGA (O-GlcNAcase) liver knockouts, or pharmacological inhibition of OGA coupled with multi-Omics analysis and bioinformatics. We identified numerous genes, proteins, phospho-proteins, or metabolites that were either inversely or equivalently changed between conditions. Moreover, we identified pathways in OGT knockout samples associated with increased aneuploidy. To test and validate these pathways, we induced liver growth in OGT knockouts by partial hepatectomy. OGT knockout livers showed a robust aneuploidy phenotype with disruptions in mitosis, nutrient sensing, protein metabolism/amino acid metabolism, stress response, and HIPPO signaling demonstrating how OGT is essential in controlling aneuploidy pathways. Moreover, these data show how a multi-Omics platform can discern how OGT can synergistically fine-tune multiple cellular pathways.

## Main Text

The fundamental challenge in understanding O-GlcNAcylation is that pharmacologic or genetic manipulation of O-GlcNAc leads to pleiotropic effects impairing data interpretation in a mechanistic way. O-GlcNAc is a ubiquitous post-translational modification found in every metazoan cell consisting of a single N-acetylglucosamine residue attached to serine or threonine residues of cytoplasmic, mitochondrial, or nuclear proteins^1^. Singular enzymes process the modification with O-GlcNAc Transferase (OGT) responsible for adding the sugar while O-GlcNAcase (OGT) removes the modification^1^. The modification is dynamic, found on thousands of proteins, and involved in regulating most cellular functions^1^. Further complicating O-GlcNAc studies, complete knockouts of OGT and OGA are lethal to dividing cells and embryos^2, 3^; while genetic or pharmacologic manipulation of one of the enzymes concomitantly changes the other enzyme^4^. These pleiotropic effects coupled to the difficulty altering the enzymes activity or expression lead to tenuous data interpretation. O-GlcNAc site mutant studies can provide mechanistic details but cannot recapitulate the complete influence of O-GlcNAcylation on cellular function. However, multi-Omic approaches coupled with genetic and pharmacologic manipulation of O-GlcNAc can provide novel information on how O-GlcNAcylation regulates cellular pathways in a synergistic manner.

Multi-Omics technologies provide mechanistic understanding of the function of O-GlcNAcylation by evaluating cellular changes at a systems level and by revealing complex relationships missed by conventional techniques. Due to the multi-faceted nature of biological processes, unimodal data will necessarily provide an incomplete mechanistic understanding. Thus, multi-Omics data captures molecular changes across multiple biological levels, offering a more holistic and robust interpretations of each “Omic” type individually. These approaches offer powerful and informative insights into biological systems; yet alone, they fail to account for the myriad interactions taking place between molecules giving rise to cellular phenotypes^5^. Molecular interaction networks provide a scaffolding through which “Omics” and interaction data can be integrated enabling the vast suite of network analysis methods to be brought to bear on “Omics” data analysis. One tool in this suite, AMEND (Active Module identification with Experimental data and Network Diffusion), is highly effective in identifying dense molecular subnetworks with large experimental values^6^. AMEND elucidates important features associated with a biological condition and the specific interactions between them. Biological network analysis techniques are especially well-suited to address the challenges presented with O-GlcNAc by providing a robust framework to tease apart the pleiotropic actions of O-GlcNAc.

To gain a more complete mechanistic understanding of O-GlcNAc regulation of cellular processes, we either knocked-out hepatocyte OGT or OGA of adult mice, or elevated O-GlcNAc for 1 to 2 weeks with OGA inhibitor Thiamet-G (TMG) ^7-9^. We performed multi-Omics and the data was analyzed using the AMEND algorithm (**Figure 1A**). We identified loss of OGT to several cellular pathways associated with aneuploidy. Previously, we identified loss of liver OGT with hyperplasia after partial liver hepatectomy (PHX) ^7, 10^; thus, we performed PHX on OGT knockout livers and found a robust increase in liver aneuploidy, validating changes in the pathways promoting aneuploidy. Together, the combination of altered O-GlcNAc animal models, multi-Omics, and advanced bioinformatics demonstrate a way to identify pathways regulated by O-GlcNAcylation that synergistically drive cellular processes preventing aneuploidy. Furthermore, this data set underscores the potential of multi-Omics to identify the mechanisms by which O-GlcNAc controls biological processes.

**Figure 1:**
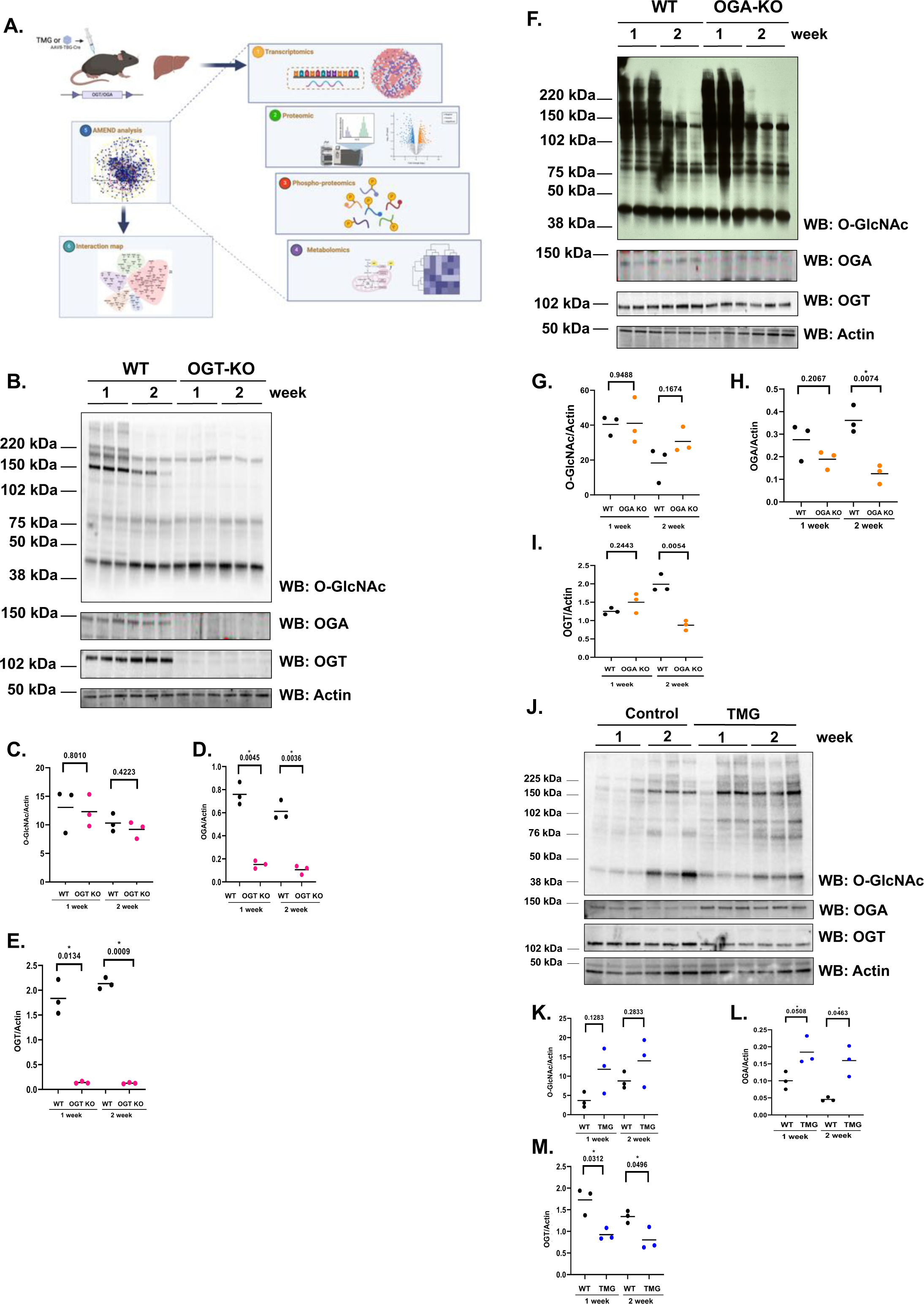
Multi-Omic based approach to understand the pleiotropic effects of O-GlcNAcylation on liver: **A.** Schematic of the multi-Omic approach: floxed-OGT or OGA mice livers were treated with AAV-cre via IP injection; alternatively, mice were treated with OGA inhibitor TMG via IP injection for 1 or 2 weeks. At 1 or 2 weeks, animals were sacrificed and livers harvest. Liver samples were used for transcriptomics, proteomics, phospho-proteomic, and metabolomics analysis. Bioinformatics was performed with the AMEND algorithm to integrate the data longitudinally or heterogeneously (Figure designed in Biorender). **B** Western blot analysis for O-GlcNAcylation, OGT, and OGA expression from OGT knockout livers at 1 and 2 weeks post-knockout. Actin is used as a load control (n=3). **C-E**: Densitometry of western blot samples. * = p value less than 0.05. **F:** Western blot analysis for O-GlcNAcylation, OGT, and OGA expression from OGA knockout livers at 1 and 2 weeks post-knockout. Actin is used as a load control (n=3). **G-I**: Densitometry of western blot samples. * = p value less than 0.05. **J:** Western blot analysis for O-GlcNAcylation, OGT, and OGA expression from mice at 1 and 2 weeks TMG treatment. Actin is used as a load control (n=3). **K-M**: Densitometry of western blot samples. * = p value less than 0.05.

## Results

### Pharmacologic and genetic manipulation of O-GlcNAc combined with multi-Omics and AMEND identifies biological pathways regulated synergistically by O-GlcNAc

Multi-Omics approaches were used to identify pathways regulated by O-GlcNAc while incorporating the pleotropic nature of O-GlcNAc into the analysis. First, we took OGT or OGA-floxed male mice and injected the animals AAV8-TBG Cre to knockout OGT or OGA in hepatocytes^7, 8, 10^, and we sacrificed the mice and harvested the livers after 1 or 2-weeks post-injection. Alternatively, we injected mice with OGA inhibitor Thiamet-G (TMG) every other day for 1 or 2-weeks and sacrificed the mice. OGT-KO liver shows a decrease in total O-GlcNAc and OGT expression in both 1 and 2 week samples (**Figure 1B-E**). OGA declines in these samples as a compensatory response^4^. OGA knockout livers showed an increase in total cellular O-GlcNAc with declines in both OGA and OGT expression (**Figure 1F-I**). TMG treated animals had increased O-GlcNAc levels and OGA expression (**Figure 1J-M**). Tissue from these animals was processed for transcriptomics, semi-quantitative proteomics (Tandem Mass Tag labeling), phospho-proteomics, or metabolomics **(Extended Tables 1A-D**). The “Omic” analysis quantified over 15,000 genes, 4500 proteins, 1500 phospho-proteins, and 250 metabolites. The resulting multi-Omics data were analyzed for differentially expressed features between treatment groups and their respective control groups. Volcano plots reveal varying coverage sizes and biological signal intensities for the different assays and group comparisons (**Extended Figure 1A-D**). Gene Set Enrichment Analysis (GSEA) was performed using log fold changes from differential expression analyses to identify dysregulated pathways (**Extended Table 1E**). We consistently identified pathways involved in the cell cycle especially mitosis, general metabolism including mitochondrial metabolism, lipid metabolism, and amino acids metabolism; as well as, pathways involved in transcription and translation. Although GSEA identified pathways targeted by O-GlcNAcylation^1^, there are two main disadvantages to the GSEA analysis. First, log fold changes involve only one treatment-control comparison, which limits our ability to generalize molecular associations to other O-GlcNAc perturbations. Second, GSEA fails to consider interactions between molecules when assessing associations between observed changes in expression and pathways. To circumvent the limitations of GSEA, we implemented the AMEND algorithm employing Equivalent Change Index (ECI), which compares log fold changes between two experiments to assess equivalent or inverse change. AMEND integrates molecular interaction information (as networks) with “Omics” data identifying densely-connected subnetworks (i.e., modules) that show coordinated dysregulation among different O-GlcNAc alterations. Next, Over-representation Analysis (ORA) was used to identify pathways showing significant overlap with AMEND modules (**Extended Table 1F**) disrupted between O-GlcNAc perturbations. Together, this analysis provides granular information on protein interactions affected by changes in O-GlcNAcylation that disrupt cellular pathways.

### Transcriptomic profiling identifies cell cycle and proteasome gene disruption in OGT KO livers

Since altered O-GlcNAc can affect the transcription of genes^11^, we profiled transcript between OGT/OGA-KO or TMG treated livers 1 and 2-week post-KO or TMG injections. OGT-KO livers between 1 and 2 weeks showed equivalently declined expression of several proteasome sub-units including proteasome-associated deubiquitinase *USP14* (**Figure 2A**), while over-representation analysis identified equivalently changed genes involved in RNA metabolism, immune system function and protein localization (**Figure 2B**). Inversely correlated genes between week 1 and week 2 of OGT-KO were predominately cell cycle genes showing substantial increases at week 2 (**Extended Figure 2A-B**). OGA-KO between 1 and 2-weeks showed equivalent changes in DNA repair and RNA Polymerase II transcription elongation (**Extended Figure 2C-D**). Inverse related changes in OGA-KO gene expression over 1 and 2-weeks included an increase in *mTOR* (Mammalian Target of Rapamycin) and genes involved in signal transduction (**Extended Figure 2E-F**). TMG 1 and 2-week treatment induced equivalent gene expression changes in RNA and glucose metabolism (**Extended Figure 2G-H**). Inversely related gene expression changes with TMG treatment include increased DNA repair genes (**Extended Figure 2I-J**). Next, we compared inverse gene expression changes in OGT-versus OGA-KO at 2-weeks post-KO. OGT-KO had elevated changes in signaling genes involved in cell cycle progression and mTORC1 signaling while OGA-KO had increased expression of genes involved in gene expression (**Figure 2C-D**). OGT- and OGA-KO showed at 2-weeks equivalently changed genes involved in extracellular matrix organization and signal transduction (**Extended Figure 3A-B**). When we compared inversely expressed genes between OGT-KO and TMG treatment at 2 weeks, we found OGT-KO increased cell cycle gene expression and decreased stress response genes (**Figure 2E-F**). Equivalently changed genes between OGT-KO and TMG included increased expression of mitotic genes, *mTOR*, and *kRAS* (**Extended Figure 3C-D**). Equivalently changed genes between OGA-KO and TMG-treatment at 2-weeks include changes in signal transduction genes and decreased expression of electron transport chain (ETC) complex 1 genes (**Figure 2G-H**). Inverse changes between OGA-KO and TMG at 2-weeks include changes in WNT (Wingless-Type MMTV Integration Site Family) signaling (**Extended Figure 3E-F**). From across the different O-GlcNAc perturbations, we consistently saw changes in mTOR signaling, cell cycle, protein metabolism, and stress response pathways.

**Figure 2:**
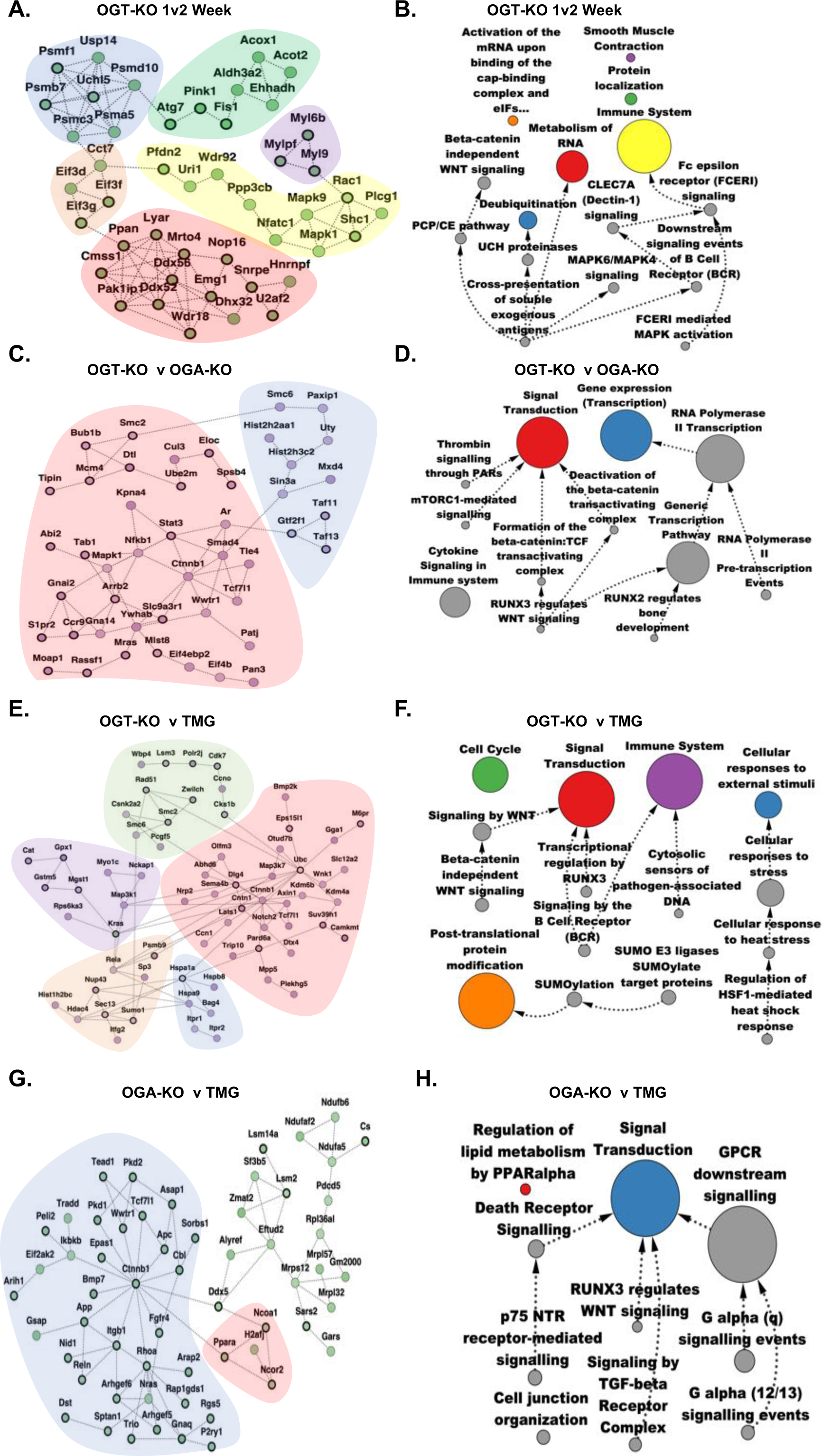
Identification of transcriptomic subnetworks using AMEND and Overrepresentation Analysis (ORA) pathways from longitudinal or heterogeneous sampling: Each AMEND subnetwork compares log fold changes between two treatment-control comparisons using the Equivalent Change Index (ECI). Green nodes represent equivalent change between treatment groups 1 and 2 with a bold border signifying that both treatment groups were up-regulated compared to their controls and a thin border signifying that both treatment groups were down-regulated compared to their controls. Purple nodes represent inverse change between treatment groups 1 and 2 with a bold border signifying that treatment 1 was up-regulated while treatment 2 was down-regulated, and vice-versa for a thin border. “T” = transcriptomic, “+” = AMEND selects for positive ECI nodes (equivalent change), “-” = AMEND selects for negative ECI nodes (inverse change). Clusters within each subnetwork are colored corresponding to the pathway associated with the nodes in that cluster. Treatment 1 and treatment 2 correspond to the first and second treatment listed in the panel description, respectively. Arrows in the ORA networks show the direction of nestedness (i.e., the source node is nested within the target node) within the directed acyclic graph and the sizes of the pathways are represented by the relative sizes of the nodes. The color of the node reflects the AMEND protein interaction network. Only the top 15 pathways, ranked by adjusted p-value, are shown. **A**: AMEND module, OGT-KO 1 week vs. 2 week (T)(+), **B**. ORA pathways, OGT-KO 1 week vs. 2 week (T)(+), **C**: AMEND module, OGT-KO vs. OGA-KO at 2 weeks (T)(-), **D**: ORA pathways, OGT-KO vs. OGA-KO at 2 weeks (T)(-), **E**: AMEND module, OGT-KO vs. TMG at 2 weeks (T)(-), **F**: ORA pathways, OGT-KO vs. TMG at 2 weeks (T)(-), **G**: AMEND module, OGA-KO vs. TMG at 2 weeks (T)(+), **H**: ORA pathways, OGA-KO vs. TMG at 2 weeks (T)(+).

**Figure 3:**
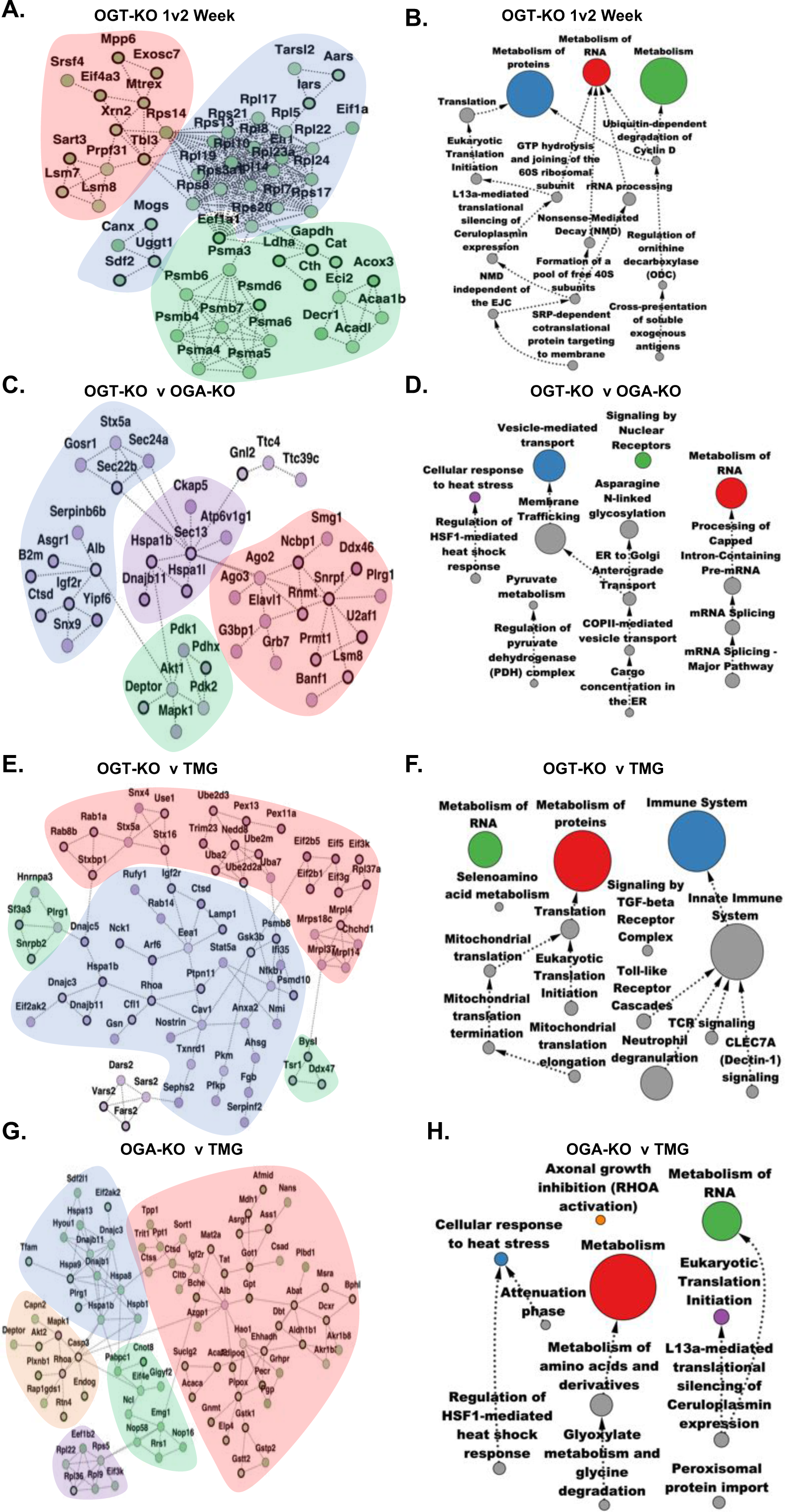
Identification of proteomic subnetworks using AMEND and Overrepresentation Analysis (ORA) pathways from longitudinal or heterogeneous sampling: Each AMEND subnetwork compares log fold changes between two treatment-control comparisons. Green nodes represent equivalent change between treatment groups 1 and 2 with a bold border signifying that both treatment groups were up-regulated compared to their controls and a thin border signifying that both treatment groups were down-regulated compared to their controls. Purple nodes represent inverse change between treatment groups 1 and 2 with a bold border signifying that treatment 1 was up-regulated while treatment 2 was down-regulated, and vice-versa for a thin border. “P” = proteomic, “+” = AMEND selects for positive ECI nodes (equivalent change), “-” = AMEND selects for negative ECI nodes (inverse change). Clusters within each subnetwork are colored corresponding to the pathway associated with the nodes in that cluster. Treatment 1 and treatment 2 correspond to the first and second treatment listed in the panel description, respectively. Arrows in the ORA networks show the direction of nestedness within the directed acyclic graph and the sizes of the pathways are represented by the relative sizes of the nodes. The color of the node reflects the AMEND protein interaction network. Only the top 15 pathways, ranked by adjusted p-value, are shown. **A**: AMEND module, OGT-KO 1 week vs. 2 week (P)(+), **B**. ORA pathways, OGT-KO 1 week vs. 2 week (P)(+), **C**: AMEND module, OGT-KO vs. OGA-KO at 2 weeks (P)(-), **D**: ORA pathways, OGT-KO vs. OGA-KO at 2 weeks (P)(-), **E**: AMEND module, OGT-KO vs. TMG at 2 weeks (P)(-), **F**: ORA pathways, OGT-KO vs. TMG at 2 weeks (P)(-), **G**: AMEND module, OGA-KO vs. TMG at 2 weeks (P)(+), **H**: ORA pathways, OGA-KO vs. TMG at 2 weeks (P)(+).

### Proteomics identifies increased stress response pathways and disrupted protein metabolism in OGT-KO liver

Since perturbations to O-GlcNAc affect protein translation and turnover^12, 13^, we performed quantitative mass spectrometry with AMEND to identify proteome changes across the different O-GlcNAc perturbations. OGT-KO livers between 1 and 2-weeks showed equivalently declined expression of several ribosome sub-units (**Figure 3A-B**) and inverse changes in pre-mRNA capping and nuclear envelop reformation (**Extended 4A-B**). OKT-KO to OGA-KO at 2-weeks demonstrated inverse changes in stress response (**Figure 3C-D**). Between OGT- and OGA-KO at 2-weeks, we found equivalent changes in the protein expression of numerous metabolic pathways (**Extended Figure 4C-D**). Inversely related protein changes between OGT-KO and TMG-treatment at 2 weeks identified protein metabolism differences between the samples (**Figure 3E-F**). Equivalent protein changes between OGT-KO and TMG-treatment at 2 weeks include lower expression of ribosomal proteins and proteasome proteins (**Extended Figure 4E-F**). Equivalent protein changes between OGA-KO and TMG-treatment at 2-weeks include changes in amino acids metabolism and decreased expression of stress response proteins (**Figure 3G-H**). Inverse change in protein expression between OGA-KO and TMG-treatment at 2 weeks include increases in ribosomal and proteasome proteins in the OGA-KO (**Extended Figure 4G-H**). Within the proteome, we again saw over-representation of protein metabolism and stress response pathways.

**Figure 4:**
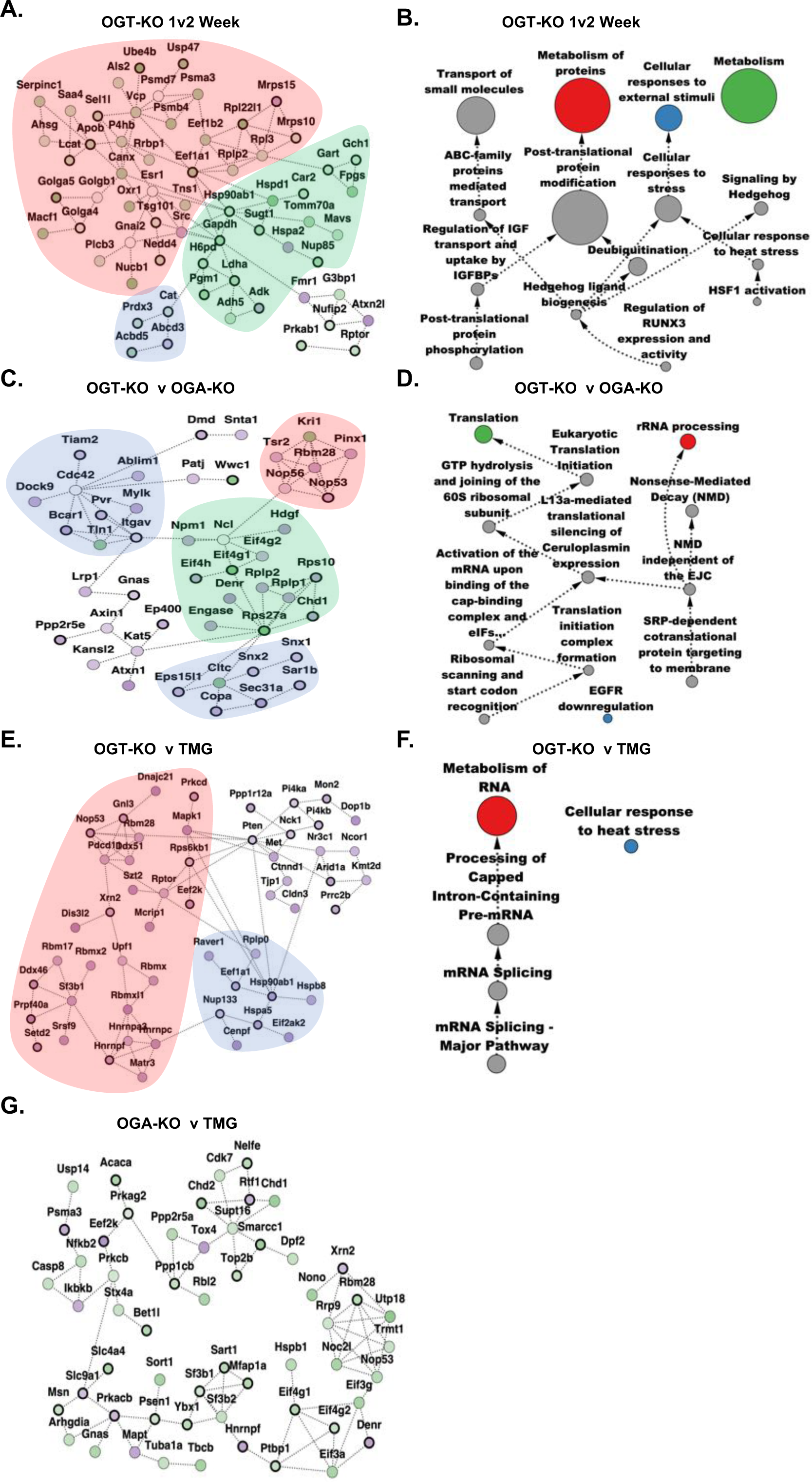
Identification of phospho-proteomic subnetworks using AMEND and Overrepresentation Analysis (ORA) pathways from longitudinal or heterogeneous sampling: Each AMEND subnetwork compares log fold changes between two treatment-control comparisons. Green nodes represent equivalent change between treatment groups 1 and 2 with a bold border signifying that both treatment groups were up-regulated compared to their controls and a thin border signifying that both treatment groups were down-regulated compared to their controls. Purple nodes represent inverse change between treatment groups 1 and 2 with a bold border signifying that treatment 1 was up-regulated while treatment 2 was down-regulated, and vice-versa for a thin border. “Ph” = phospho-proteomic, “+” = AMEND selects for positive ECI nodes (equivalent change), “-” = AMEND selects for negative ECI nodes (inverse change). Clusters within each subnetwork are colored corresponding to the pathway associated with the nodes in that cluster. Treatment 1 and treatment 2 correspond to the first and second treatment listed in the panel description, respectively. Arrows in the ORA networks show the direction of nestedness within the directed acyclic graph and the sizes of the pathways are represented by the relative sizes of the nodes. The color of the node reflects the AMEND protein interaction network. Only the top 15 pathways, ranked by adjusted p-value, are shown. **A**: AMEND module, OGT-KO 1 week vs. 2 week (Ph)(+), **B**. ORA pathways, OGT-KO 1 week vs. 2 week (Ph)(+), **C**: AMEND module, OGT-KO vs. OGA-KO at 2 weeks (Ph)(-), **D**: ORA pathways, OGT-KO vs. OGA-KO at 2 weeks (Ph)(-), **E**: AMEND module, OGT-KO vs. TMG at 2 weeks (Ph)(-), **F**: ORA pathways, OGT-KO vs. TMG at 2 weeks (Ph)(-), **G**: AMEND module, OGA-KO vs. TMG at 2 weeks (Ph)(+), **H**: ORA pathways, OGA-KO vs. TMG at 2 weeks (Ph)(+).

### Phospho-proteomics identifies alterations to ribosome and heat shock pathway phosphorylation in OGT-KO livers

There is a strong relationship between changes in O-GlcNAcylation affecting phosphorylation^14^; thus, we used quantitative proteomics to measure the impact of O-GlcNAc perturbations on the phospho-proteome. After phospho-peptide enrichment and proteomics, AMEND analysis found equivalently changed phosphorylations of ribosome, proteasome, and ubiquitinylation complex proteins and increased phosphorylation of HSP90 (heat shock protein 90) in OGT-KO livers between 1 and 2-weeks, (**Figure 4A-B**). Inverse phosphorylation changes between OGT-KO at 1 and 2-weeks include alterations to protein metabolism, RNA metabolism, and heat shock pathways (**Extended 5A-B**). Inverse phosphorylation changes between OGT-KO and OGA-KO at 2-weeks include protein translation and rRNA processing pathways (**Figure 4C-D**). Equivalent phosphorylation changes between OGT-KO and OGA-KO at 2-weeks include signal transduction pathways and metabolism of RNA (**Extended 5C-D**). Inverse phosphorylation changes between OGT-KO and TMG treatment at 2-weeks include changes in RNA metabolism and response to heat stress pathways (**Figure 4F-E**). Equivalent changes between OGT-KO and TMG treatment at 2-weeks were seen in RNA metabolism (**Extended Figure 5F-E**). Equivalent phosphorylation changes between OGA-KO and TMG at 2-weeks showed decreased phosphorylation of USP14 (**Figure 4G**). However, no pathway was over-represented in the comparison. Inverse phosphorylation changes between OGA-KO and TMG treatment at 2-weeks include pathways involved in signal transduction, RNA metabolism, and the cell cycle (**Extended Figure 5G-H**). Over-represented pathways in the phospho-proteomics again showed O-GlcNAc perturbations affecting the cell cycle, protein metabolism, and stress response

**Figure 5:**
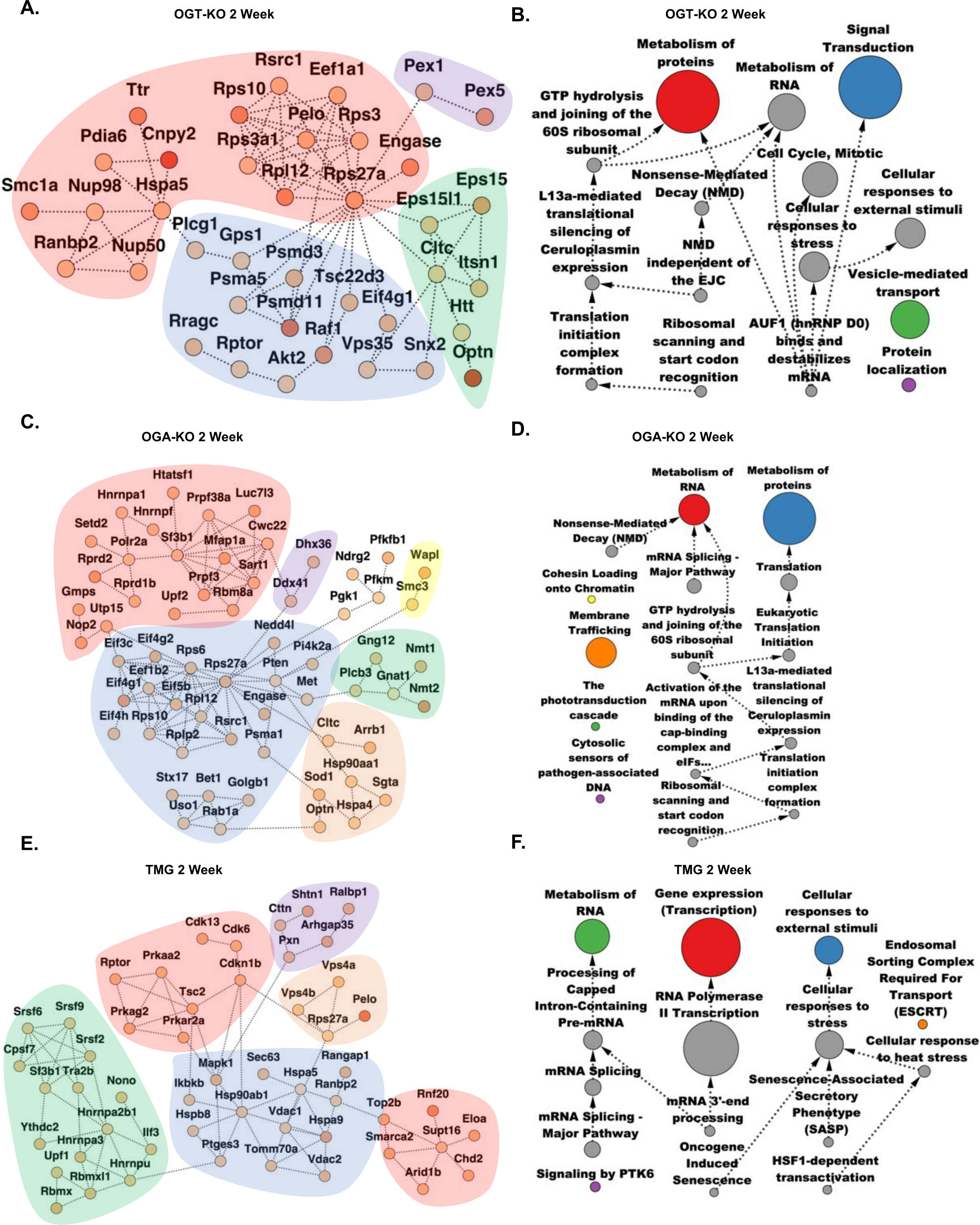
Differential phosphorylation subnetworks from AMEND, along with pathways from Overrepresentation Analysis (ORA) represented as a directed acyclic graph: Each AMEND subnetwork searched for nodes with large absolute differences in log fold change between proteomic and phosphoproteomic data for a given treatment-control comparison computed as a Welch’s t-test statistic. Darker shades of orange represent larger t-test statistics. Clusters within each subnetwork are colored corresponding to the pathway associated with the nodes in that cluster. Arrows in the ORA networks show the direction of nestedness (i.e., the source node is nested within the target node) and the sizes of the pathways are represented by the relative sizes of the nodes. Only the top 15 pathways, ranked by adjusted p-value, are shown. **A**: AMEND module, OGT-KO at 2 weeks, **B**: ORA pathways, OGT-KO at 2 weeks, **C**: AMEND module, OGA-KO at 2 weeks, **D**: ORA pathways, OGA-KO at 2 weeks, **E**: AMEND module, TMG at 2 weeks, **F**: ORA pathways, TMG at 2 weeks.

Next, we looked at large phosphorylation changes in specific proteins. In OGT-KO 2-week livers, we measured robust phosphorylation changes in pathways involved in protein metabolism, cell cycle, and signal transduction (**Figure 5A-B**). In OGA-KO 2-week livers we measured phosphorylation changes in pathways linked to RNA metabolism and protein metabolism (**Figure 5C-D**). Finally, we measured robust phosphorylation changes in 2-week TMG treated livers in pathways linked to RNA metabolism and gene transcription (**Figure 5E-F**). These data again point to cell cycle related pathways altered by perturbed O-GlcNAcylation.

### Metabolomics identifies OGT regulation of glycine, serine, and threonine metabolism

Due to known O-GlcNAc perturbations affecting metabolic processes^8, 10^, we performed discovery based proteomics and AMEND analysis to determine metabolic pathways affected by changes in O-GlcNAcylation. Metabolic pathways equivalently changed in OGT-KO livers between 1 and 2-weeks include increases in pathways involving glycine, serine, and threonine metabolism (**Figure 6A-B**). Inversely changed metabolic pathways from OGT-KO 1 to 2-week include methionine and propanoate metabolism (**Extended Figure 6A-B**). Equivalently changed metabolic pathways in OGA-KO from 1 to 2-weeks include increases in glycine and glutamate (**Extended Figure 6C-D**). Inversely changed metabolic pathways from OGT-KO 1 to 2-week include changes in arginine and proline metabolism (**Extended Figure 6E-F**). Inversely correlated metabolic pathways between OGT- and OGA-KO at 2-weeks include increased glutathione metabolism in the OGT-KO (**Figure 6C-D**). Equivalently changed metabolic pathways between OGT- and OGA-KO at 2-weeks include increased glycine metabolism (**Extended Figure 6G-H**). Inversely changed metabolites between OGT-KO and TMG at 2-weeks include increased glycine and methionine in OGT-KO (**Figure 6E-F**). Equivalently changed metabolites between OGT-KO and TMG at 2-weeks include glucosamine-6-phosphate (**Extended Figure 6I-J**). Equivalently changed metabolites between OGA-KO and TMG at 2-weeks are reduced levels of glucosamine and glucosamine-6-phosphate (**Figure 6G**), which are precursor metabolites for UDP-GlcNAc, the metabolic substrate for OGT, and inversely correlated pathways include amino acid metabolism (**Extended Figure 6K-L**). In summary, we consistently saw changes in glycine, serine, and threonine metabolism with the O-GlcNAc perturbations.

**Figure 6:**
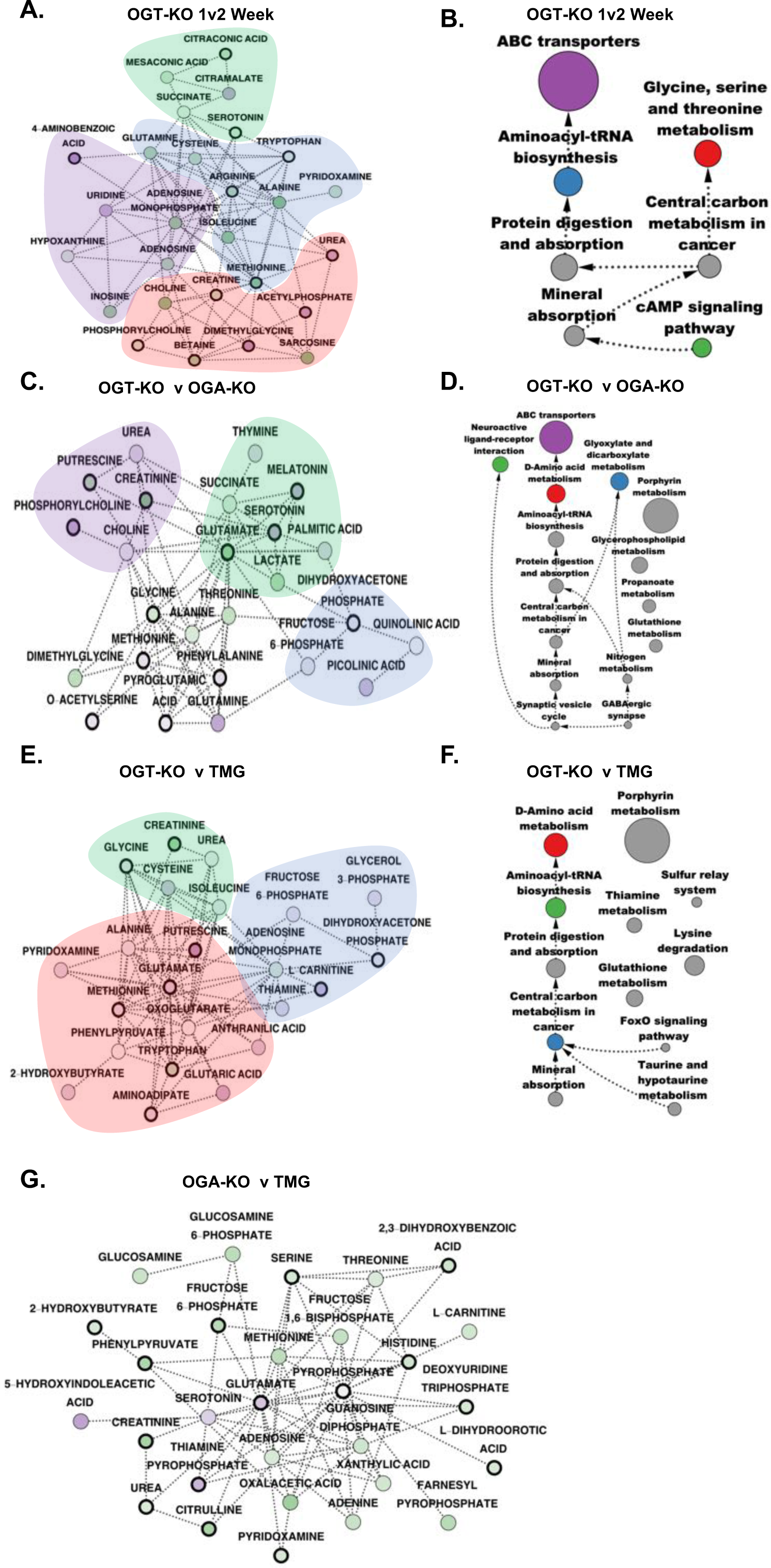
Identification of metabolomic subnetworks from using AMEND and Overrepresentation Analysis (ORA) represented as a directed acyclic graph from longitudinal or heterogeneous sampling: “M” = metabolomic, “+” = AMEND selects for positive ECI nodes (equivalent change), “-” = AMEND selects for negative ECI nodes (inverse change). Clusters within each subnetwork are colored corresponding to the pathway associated with the nodes in that cluster. Treatment 1 and treatment 2 correspond to the first and second treatment listed in the panel description, respectively. Arrows in the ORA networks show the direction of nestedness (i.e., the source node is nested within the target node) and the sizes of the pathways are represented by the relative sizes of the nodes. Only the top 15 pathways, ranked by p-value, are shown. **A**: AMEND module, OGT-KO 1 week vs. 2 week (M)(+), **B**: ORA pathways, OGT-KO 1 week vs. 2 week (M)(+), **C**: AMEND module, OGT-KO vs. OGA-KO at 2 weeks (M)(-), **D**: ORA pathways, OGT-KO vs. OGA-KO at 2 weeks (M)(-), **E**: AMEND module, OGT-KO vs. TMG at 2 weeks (M)(-), **F**: ORA pathways, OGT-KO vs. TMG at 2 weeks (M)(-), **G**: AMEND module, OGA-KO vs. TMG at 2 weeks (M)(+). No pathways were found to be significantly associated with the OGA-KO vs. TMG subnetwork.

### Liver hepatectomy in OGT-KO mice increased aneuploidy as predicted by AMEND

After AMEND analysis, we identified the following pathways over-represented in OGT-KO analysis: cell cycle, HIPPO signaling, mTOR/AMPK signaling, p53 function, proteasome function, stress response, and serine flux. Next, we looked for biological functions all these pathways have in common, and we identified aberrant function of these pathways is linked to aneuploidy^15-24^. Moreover, Hepatocytes can be polyploid, and hepatocyte aneuploidy increases with aging^25^. Thus, to test the potential OGT driven increase in aneuploidy, we measured DNA content in OGT-KO livers. The OGT-KO livers showed some polyploidy but no aneuploidy (**Figure 7A**). Previously, when OGT-KO livers were subjected to partial liver hepatectomy (PHX), the livers showed increased cell growth and hypertrophy after 1-month^7, 10^; therefore, we challenged control and OGT/OGA-KO livers by performing a PHX and collecting the tissue 2-weeks post PHX. After PHX, OGT-KO livers presented with a robust aneuploid phenotype as measured by DNA content (**Figure 7B**). Furthermore, OGA-KO livers were phenotypically like control (**Extended Figure 7A**). OGA-KO livers did not have over-representation of the pathways seen in the OGT-KO and serves as a negative control for PHX after O-GlcNAc disruption.

**Figure 7:**
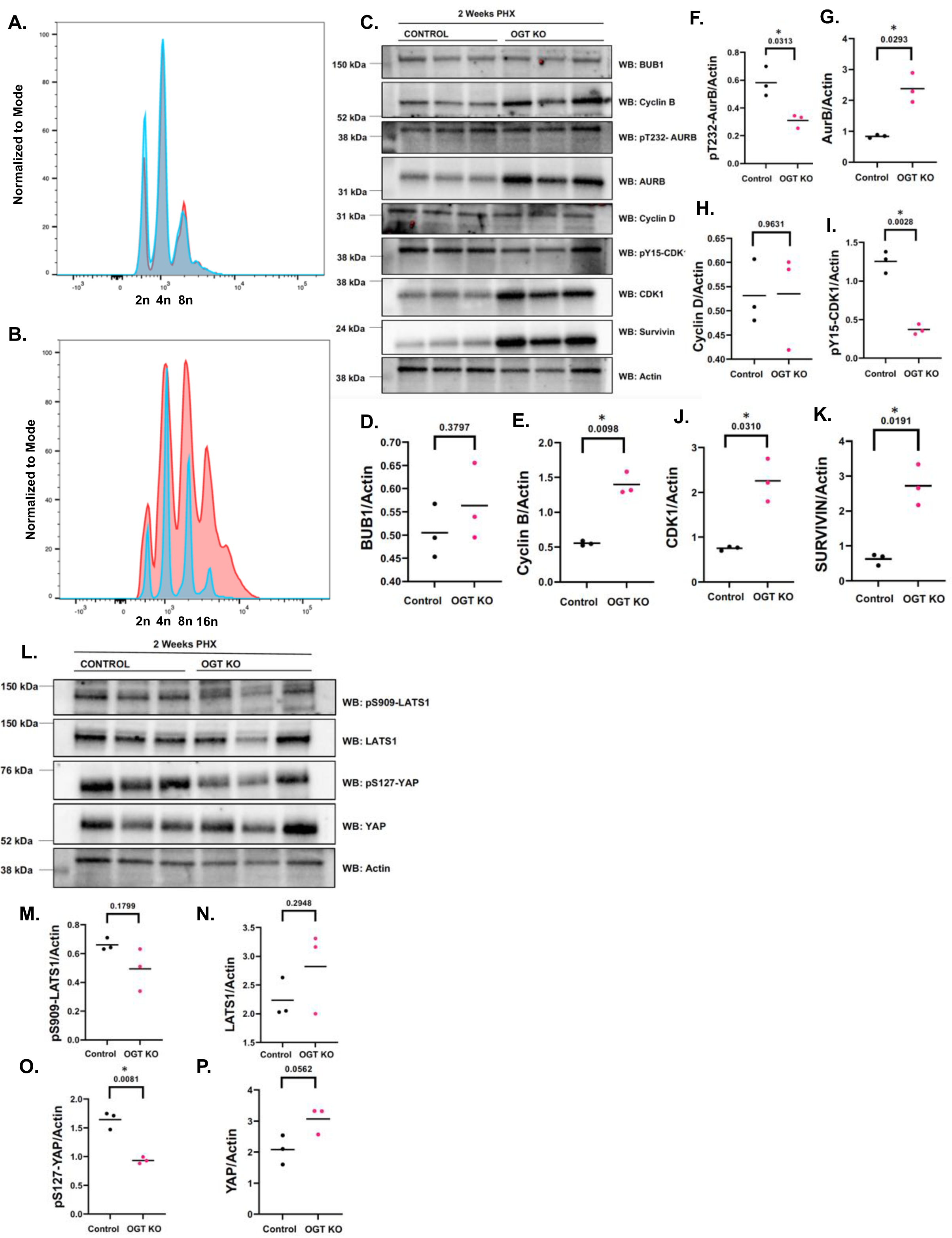
OGT KO Livers have increased aneuploidy after hepatectomy: **A:** Flow cytometry was performed on fixed, propidium iodine stained livers. Control livers after PHX were polyploid (blue is no PHX, red is after PHX) **B.** OGT KO livers had greater then 4n and aberrant DNA content after PHX (blue is no PHX, red is after PHX). **C:** Western blot analysis on mitotic proteins. Actin is used as a load control (n=3). **D-K:** Densitometry of western blot samples. * = p value less than 0.05. **L:** Western blot of HIPPO signaling pathway proteins (n=3). Actin is used as a load control **M-P:** Densitometry of western blot samples. * = p value less than 0.05.

Next, we validated the expression and phosphorylation status of mitotic proteins associated with aneuploidy. Consistent with the over-represented pathways, we found significant increases in the expression of Cyclin B, Cyclin Dependent Kinase 1 (CDK1), Aurora Kinase B (AurB), and AurB interacting protein Survivin (**Figure 7C-K**). Moreover, we found decreased levels of inhibitory Tyrosine 15 phosphorylation on CDK1 and decreased activating phosphorylation of T232 on AurB (**Figure 7C-K**). Importantly, loss of AurB T232 phosphorylation promotes aneuploidy^26^. OGA-KO PHX livers showed no significant changes in mitotic protein expression or phosphorylation (**Extended Figure 7B-J**). This result supports the critical role of OGT in maintaining the homeostasis of mitotic protein function through O-GlcNAcylation^16^.

AMEND identified changes in HIPPO signaling proteins in OGT-KO livers, and increased HIPPO signaling promotes aneuploidy^17^. We measured the expression and phosphorylation status of HIPPO downstream kinase LATS1 (Large Tumor Suppressor Kinase 1) and transcriptional activator YAP (Yes-associated protein). After PHX, we found a significant decrease in YAP inhibitory phosphorylation by LATS1 (**Figure 7L-P**). OGA-KO PHX livers showed no significant changes in HIPPO signaling (**Extended Figure 7K-O**).

Moreover, AMEND connected transcriptome, proteome, and phospho-proteome changes in mTOR/AMPK signaling in OGT-KO livers to potential induction of aneuploidy^18, 19^. We measured mTOR signaling pathway proteins in OGT-KO livers after PHX. Although we found a significant decrease in mTOR and S6K expression, mTOR activating phosphorylation of S6 Kinase (S6K) was increased. Furthermore, we found an increase in expression of Oxysterol Binding Protein (OSBP), an activator of mTOR^27^ whose over-expression promotes aneuploidy (**Figure 8A-H**)^28^. Interestingly, OGA-KO liver PHX showed an increase in S6K phosphorylation but little to no change in S6 phosphorylation or expression (**Extended Figure 7P-W**). Subsequently, we analyzed AMP-regulated kinase (AMPK) function, which inhibits mTOR signaling during nutrient stress^29^. We found that total AMPK protein levels were significantly decreased in OGT-KO livers, and AMPK substrate ACC1 (Acetyl-CoA Carboxylase 1) phosphorylation was unchanged (**Figure 8I-M**). We measured no changes in AMPK signaling in the OGA-KO livers after PHX (**Extended Figure 7X-BB**).

**Figure 8:**
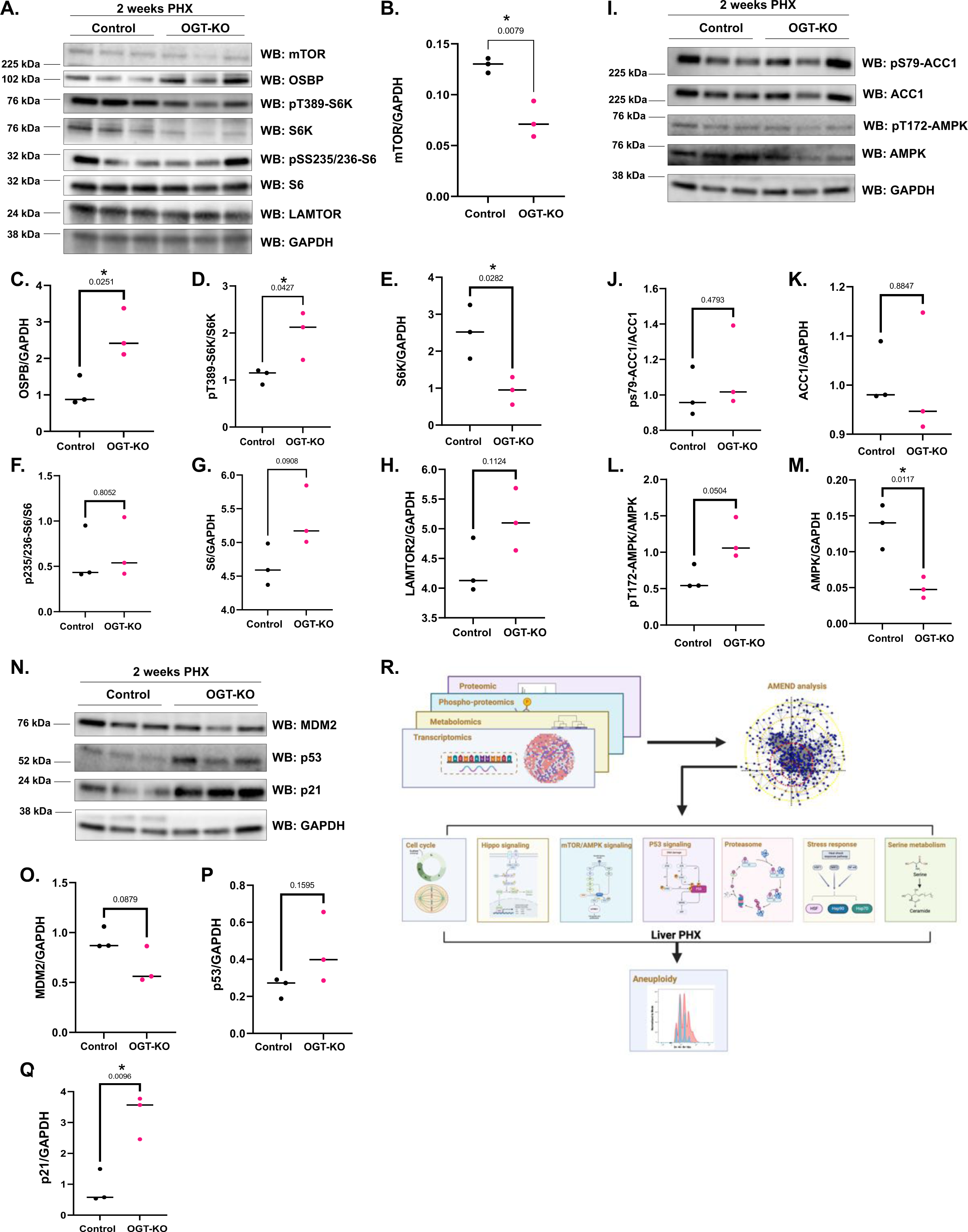
Loss of OGT activates several pathways involved in aneuploidy: **A:** Western blot of mTOR signaling pathway proteins (n=3). GAPDH is used as a load control (n=3). **B-H:** Densitometry of western blot samples. * = p value less than 0.05. **I:** Western blot of AMPK signaling pathway proteins (n=3). GAPDH is used as a load control (n=3). **J-M:** Densitometry of western blot samples. * = p value less than 0.05. **N:** Western blot of p53 signaling pathway proteins (n=3). GAPDH is used as a load control (n=3). **O-Q:** Densitometry of western blot samples. * = p value less than 0.05. **R:** Summary: Multi-Omic approached coupled with genetic and pharmacologic manipulation of O-GlcNAc identified several pathways linked to increased aneuploidy changed in OGT KO livers. Flow cytometry of OGT KO livers after PHX showed increased aneuploidy and validation western blots displayed changes consistent with aneuploidy (Figure designed in Biorender).

Lastly, robust p53 (Tumor Suppressor Protein 53) activation leads to cell cycle arrest and apoptosis; however, mitotic defects associated with AurB function activate a subset of p53 target genes characterized by induction of cell cycle inhibitor protein p21 (CDK inhibitor p21) to promote aneuploidy^30^. We measured p53 signaling after OGT-KO liver PHX at 2-weeks and saw no changes in the expression of p53 or p53-regulator protein MDM2 (Double minute 2), but p21 levels were significantly increased (**Figure 8N-Q**). OGA-KO PHX liver at 2-weeks showed no changes in p53 or MDM2, but p21 levels were increased (**Extended Figure 7CC-FF**). Together, these data validate the over-represented pathways identified in the multi-Omic analysis.

## Discussion

Herein, we demonstrate that a multi-Omics approach combined with AMEND analysis circumvents inherent problems and pleotropic effects with O-GlcNAc manipulation revealing that OGT expression is essential for preventing aneuploidy by synergistically regulating pathways contributing to the development of liver aneuploidy (**Figure 8R)**. We identified multiple cellular pathways disrupted by loss of OGT that contribute to aneuploidy including cell cycle/mitosis, Hippo Signaling, mTOR/AMPK, p53, Proteasome, Stress Response, and serine metabolism. Critically, all these pathways perform essential cellular functions that need to be tightly controlled for cell health; O-GlcNAcylation provides the cell a dynamic and responsive rheostat to quickly modulate all these pathways to maintain cellular homeostasis.

The Cell cycle is an excellent example of O-GlcNAc homeostatic regulation. Over-expression of OGT and OGA delays mitotic exit while OGT over-expression increases chromosomal instability resulting in aneuploid cells^31, 32^. OGT is strongly associated with both the mitotic and meiotic spindles^31, 33^, spindles are highly O-GlcNAcylated^16^, and both OGT and OGA move to the mid-body during cytokinesis via an interaction with Aurora Kinase B^34^. The rate of O-GlcNAc addition and removal, termed O-GlcNAc cycling, is critical for spindle organization since OGT/OGA over-expression both cause an increase in the length and width of the mitotic spindle, while TMG reduced length and width^8^. The dynamics of O-GlcNAc cycling appear critical for organizing mitotic progression since over-expression of either OGT or OGA cause the same spindle phenotype. We hypothesize this is due to increased cycling that disrupts the ability of protein complexes to organize properly since over-expression of either OGT or OGA causes a concurrent change in the other enzyme resulting in a net increase in the rate of cycling^8^. TMG slows cycling, allowing for protein complexes to organize correctly. Thus, the on/off rate of O-GlcNAc allows the modification to act as a rheostat to fine-tune protein function across pathways.

All nutrient sensors are rheostats that modulate cell function in response to nutrient changes. For example, mTOR signaling senses branch chain amino acids and growth factors promoting cell growth while AMPK senses AMP levels slowing growth and counter-balancing mTOR^29^. O-GlcNAcylation has strong synergy with these pathways. AMPK can phosphorylate OGT while also being a substrate for OGT with increased AMPK O-GlcNAcylation correlating with increased activity^35, 36^. Moreover, higher O-GlcNAcylation is associated with decreased mTOR signaling^36^ while OGT loss induces mTOR activation^37^. Our data showing lower AMPK levels and decreased AMPK signaling with higher mTOR signaling corroborates these findings. Our approach further argues for the idea that OGT is a nutrient sensing hub. First, the concentration of the metabolic substrate for OGT, UDP-GlcNAc (Uridine-Diphosphate N-acetyl-glucosamine), is sensitive to changes in carbohydrates, lipids, nucleotides, and amino acids^38^ linking OGT function to the nutrient environment. Second, there is only one OGT gene, making OGT the control point for integrating nutrient signals^2^. Third, OGT activity increases as availability of UDP-GlcNAc increases^39^; and fourth, most cellular pathways contain O-GlcNAcylated proteins^40^ allowing OGT to regulate most cellular functions according to nutrient levels. OGT as nutrient gatekeeper explains why OGT knockout is lethal in embryonic tissue and why loss of OGT can impact diverse aneuploidy associated pathways.

We identified 7 pathways linked to aneuploidy development. Besides mitotic pathways and nutrient sensors, the p53 pathway and stress response pathways are well-known targets of O-GlcNAcylation. Aneuploidy in liver is not uncommon after a stress induced injury^25, 41^, and we measured alterations of stress-response pathways linked to aneuploidy^22^. Loss of p53 is associated with aneuploidy^20^, and although we did not see a loss in p53 or MDM2 expression, p53 target p21 was up-regulated with OGT-KO, and increased p21 is associated with aneuploidy involving AurB alterations^30^. We also measured changes in HIPPO signaling specifically decreases in YAP phosphorylation leading to more YAP target gene expression^7^. Conversely, YAP O-GlcNAcylation is correlated with higher activity and OGT knockdown reduces this activity^42^. Our results are opposite to these but could be due to the actions of YAP in liver, which drives quiescent hepatocytes to replicate upon liver damage^43^. Additionally, we identified changes in numerous proteasome/ubiquitin ligase proteins linked to aneuploidy including decreased USP14 expression and increased phosphorylation of NEDD4^21, 23, 44^. Lastly, serine metabolism is critical for aneuploid cell survival due to increased sphingolipid synthesis^24^; O-GlcNAc is a key regulator of serine metabolism^45^, and we measured altered serine metabolites in the OGT-KO cells.

In conclusion, we demonstrated, using multi-Omics combined with novel bioinformatics and O-GlcNAc manipulation, the essential role O-GlcNAc plays in organizing pathways associated with aneuploidy. Broadly, these data show how O-GlcNAc is a nutrient sensing rheostat and demonstrates the essential role OGT plays in integrating these signals as a singular control point within the cell to modulate pathway function.

## On-line methods

### Animal protocols and models

The University of Kansas Medical Center Animal Care and Use Committee approved all experiments in this study. Two-months old male C57Bl/6J mice and floxed-OGT mice were purchased from The Jackson Laboratory (Bar Harbor, ME). OGA floxed mice were provided by John Hanover (NIH). All mice were housed using a standard 12-h light/dark cycle with access to chow and water *ad libitum*. Mice were treated with a 50 mg/kg thiamet-G intraparietal injection every other day for 1 or 2 weeks^46^. After TMG completion, mice were fasted for 16 h before isoflurane (Fisher) anesthesia assisted cervical dislocation. Two-month-old OGT-floxed and OGA-floxed mice were injected intraperitoneally with cre recombinase driven by thyroxin binding globulin promoter carried by the AAV8 virus (AAV8-TBG-CRE) from Vector Biolabs at a 2.5E^11^ virus particles dissolved in 300 μL of saline to generate hepatocyte-specific OGT knockout (OGT-KO) and OGA knockout (OGA-KO), respectively. OGT-floxed and OGA-floxed mice treated with an AAV8 carrying GFP (AAV8-TBG-GFP) were used as controls. After one week of washout period, OGT floxed, OGT-KO, OGA-floxed, and OGA-KO mice were subjected to two-third partial hepatectomy (PHX), as previously described^47^. Mice were euthanized at 14-days post PHX, and blood and liver tissue samples were collected for further analysis.

### Antibodies

We used primary and secondary antibodies at 1:2,000 and 1:10,000 dilutions, respectively. The following antibodies were used for immunoblotting: OGT (AL-34) and OGA (345) were generously provided by Dr. Gerald Hart (Department of Biochemistry and Molecular Biology, the University of Georgia); anti-O-linked N-acetylglucosamine antibody (RL2, Thermo Fisher, MA1-072); anti-Actin antibody (Sigma Aldrich, A2066); Survivin (6E4, Cell Signaling, 2802S); p-Aurora A/B/C (T288, T232, T198) (D13A11, Cell Signaling, 2914P); monoclonal anti-AURKB antibody (Sigma Aldrich, WH0009212M3); p-cdc2 (Tyr15) (10A11, Cell Signaling, 4539S); anti-CDK1 antibody (A17, Abcam, ab18); anti-BUB1 antibody (EPR18947, Abcam, ab195268); anti-Cyclin B1 antibody (Sigma Aldrich, C8831); anti-Cyclin D1 (Sigma Aldrich, SAB4502602); phospho-YAP antibody (Ser127) (Cell Signaling, 4911); YAP (D8H1X, Cell Signaling, 14074); phospho-LATS1 (Ser909) (Cell Signaling, 9157); LATS1 (C55B5, Cell Signaling, 3477; Phospho-p70 S6 Kinase (Thr389) (Cell signaling #9205); p70 S6 Kinase Antibody (Cell signaling #9202). OSBP (Proteintech #11096-1-AP); MDM2 (Proteintech #27883-1-AP); AMPK-alpha (Cell signaling #5831S); p-AMPK (T172) (Cell signaling #2535S); p53 (Santa Cruz #G3014); p21 (Santa Cruz #A2413); ACC1(Cell signaling #4190S); p.ACC1 (Ser79) (Cell signaling#11818S); phospho-S6 (S235/236) (Cell signaling#4858S); S6 (S235/236) (Cell signaling#2217S); mTOR (Cell signaling#2972S); GAPDH (abcam #ab9485); goat anti-rabbit IgG-HRP (Bio-Rad, 170-6515; goat anti-mouse IgG-HRP (Bio Rad, 170-6516); and anti-chicken IgY-HRP (Sigma Aldrich, A9046).

### Cell Lysis and Immunoblotting

We lysed the animal tissue using RIPA buffer, which was prepared by adding the following components: 10 mM Tris (pH 7.6), 150 mM NaCl, 40 mM GlcNac, 2 mM EDTA, 1mM DTT, 1% Nonidet P-40, 0.1% SDS, 0.5% deoxycholic acid with the phosphatase inhibitors beta-glycerophosphate (1mM), 1mM sodium fluoride (NaF, 1mM) and protease inhibitors 2mM phenylmethylsulphonyl fluoride (PMSF, 2 mM). 1X inhibitor cocktail composed of 1 ug/mL leupeptin, 1 ug/mL antipain, 10 ug/mL benzamidine, and 0.1% aprotinin was added right before cell lysis. Liver tissue was homogenized by using a dounce homogenizer for at least 10 strokes or until tissue was completely homogenized. The cell lysates were left on ice for 20 minutes and vortexed every 5 minutes; they were centrifuged at 15,000 x g for 20 minutes at 4 °C to obtain homogenous proteins. The protein concentration was determined by using Bradford assay (Bio-Rad Catalog). We added 4X protein solubility mixture that includes: 100 mM Tris [pH 6.8], 10 mM EDTA, 8% SDS, 50% sucrose, 5% beta-mercaptoethanol, and 0.08% Pyronin Y, to denature the proteins. Protein lysates were activated by boiling for five minutes at 95 °C. Equal amount of protein samples was loaded on 4-15% Criterion precast TGX gels (Bio-Rad). The gels were run at constant 125 V for one hour; they then got transferred to polyvinylidene difluoride (PVDF) membranes at constant 0.4 A. Membranes were blocked with 3% BSA and 0.01% sodium azide (NaN_3_) in 1X TBST (25 mM Tris [pH 7.6], 150 mM NaCl, and 0.05% Tween-20) for 30 minutes. Blots were incubated with specific primary antibodies (1: 2,000 dilution) overnight at 4 °C. The next day, blots were washed three times in 1X TBST, 10 minutes each time. They were incubated for one hour at room temperature by using HRP-conjugated secondary antibodies in 1: 10,000 dilutions (Bio-Rad). The washing step was repeated (three times in 1X TBST, 10 minutes each) and immunoblots were developed using chemiluminescence HRP-antibody detection method (Thermo Fish, 34095). Blots were stripped in Restore PLUS Western Blot Stripping Buffer (Thermo Fish, 46430) for 20 minutes at room temperature; they were thoroughly washed using ddH_2_O and three times in 1X TBST, five minutes each time, blocked with TBST 3% BSA for 20 minutes before re-probing with a different primary antibody. Protein quantification data was obtained by using either ImageJ 3.2 (National Institutes of Health) or Image Lab (Bio-Rad); the band density of all proteins in this study was measured, from which we calculated the ratios of the protein of interest against the control protein, such as GAPDH or actin. We repeated three independent experiments for immunoblotting.

### Transcriptomic experimental preparation

The cDNA library was prepared using Illumina TruSeq Stranded mRNA sample preparation kit (Illumina) following the manufacturer’s instruction. Total RNA was isolated using the same method as described previously^8^, and total RNA per reaction was 800 ng. Samples were analyzed by the Genomics Core Facility at KUMC using Illumina NovaSeq 6000 Sequencing System as described previously^10^.

### Proteomics experimental preparation

#### Proteolysis and TMT labeling

Trypsin digestion was performed on lysates following the manufacture’s instruction included in the Tandem Mass Tag labeling kits (ThermoFisher Scientific), Briefly, 100 μg of protein per condition were lyophilized and resuspended in 100 µl of 100 mM triethylammonium bicarbonate buffer (TEAB). Following reduction and alkylation the proteins were precipitated by the addition of 6 volumes of chilled acetone and the tubes were kept at -20°C for 18h. After centrifugation, the protein pellets were resuspended in 100 ul of 50 mM TEAB and 2.5 µg of trypsin (Promega) were added per 100μg of protein and incubated at 37°C overnight.

Tandem Mass Tag labeling of the individual samples was done after digestion^48^. Briefly, 41 μl of the TMT label reagent was added to each 100μg of digested protein sample and incubated at room temperature for an hour. Then 8 μl of 5% hydroxylamine were added to each tube and incubated 15 min at room temperature to quench the reaction. Two labeling protocols were used: a six-plex (TMTsixplex™ ThermoFisher Scientific), and a 16-plex (TMTpro™ 16plex, ThermoFi7sher Scientific) The samples (in triplicate) were labeled as follows: a) for the 6plex: 126, Control; 127, TMG treated; 128 wild type; 129, OGT KO; 130, 14 days KO; 14 days, wild type. And b) for the 16-plex: 126C, OGT KO 1;127N OGT KO 2; 127C, OGT KO 3; 128N, OGT control 1; 128C OGT control 2; 129N, OGT control 3; 129C, OGA control 1; 130N, OGA control 2; 130C, OGA KO 1; 131N, OGA KO 2;131C, OGA KO 3; 132N, TMT treated 1; 132C, TMT treated 2; 133N, TMT treated 3; 133C, Saline 1; and 134C, Saline 2. Following labeling, equal amounts of each labeled sample were combined, lyophilized and stored at -80°C until fractionation and LC MS analysis.

#### High pH off-line fractionation

To increase the number of proteins identified, the combined samples were fractionated offline using high pH reversed phase chromatography. For the 6plex experiment, fractionation was done using the Pierce™ High pH Reversed-Phase Peptide Fractionation Kit (ThermoFisher Scientific); and for the 16plex offline fractionation was performed^49^. Briefly, the lyophilized sample was resuspended in 600ul 4.5 mM ammonium formate, 2% ACN, pH 10 (solvent A) and loaded into a reverse phase column (Agilent 300 Extend-C18 5 um, 4.6 x 250 mm) previously conditioned and equilibrated with solvent A. The peptides were eluted with a gradient of solvent B (4.5 mM ammonium formate, 90% CAN, pH 10) into solvent A using the following gradient: 0% B for 7 min; 0 to 16% B in 6 min; 16 to 40% B in 60 min; 40 to 44% B in 4 min; 44 to 60% B in 5 min; 60% B for 14 min. The flow rate was set at 1 ml/min. A total of 96 fractions were collected. The experiment was performed in duplicate, and the eluted fractions were combined into a total of 12 or 24 fractions, respectively. The combined fractions were lyophilized and stored at -80°C until LC MS analysis.

#### Phosphopeptide enrichment

For phosphopeptide enrichment of the 16-plex experiment, a sequential enrichment of metal oxide affinity chromatography (SMOAC) protocol was used with minor modifications^50^. The offline fractions were reduced to 6 fractions by combining every 5^th^ fraction The protocol involves the sequential use of two consecutive chromatographic steps. First, the phosphopetides in the sample are enriched using a titanium dioxide affinity chromatography (ThermoFisher Scientific High-Select TM TiO 2 phosphopeptide enrichment kit). The phosphopeptides in the unbound eluate are further enriched by ferric nitroacetate affinity chromatography (ThermoFisher Scientific High-Select TM Fe-NTA phosphopeptide enrichment kit). All steps of enrichment were done following the manufacture’s protocol.

#### Mass spectrometric analysis

Peptide fractions from the high pH fractionation were analyzed by on line by LC MS, using a UHPLC system (nLC 1200, ThermoFisher Scientific) connected on line to an Orbitrap Fusion Lumos mass spectrometer (ThermoFisher Scientific). In brief, all fractions were dissolved to a final volume of 50 (proteomes) or 20 (phospho enriched peptides) µl of 0.1% formic acid and 10 μl were direct loaded on a C18 reverse phase peptide trap column (Acclaim PepMap 75 um x 2cm ThermoFisher Scientific). Following a wash with 0.1% formic acid for 15 minutes at 0.5 µl/min, the flow of the peptide trap was diverted to the resolving column (Acclaim PepMap 75 um x 50 cm) and the peptides were eluted with the following gradients of 80% acetonitrile, 0.1 % formic acid (solvent B) into 3% acetonitrile, 0.1% formic acid (solvent A) at a flow rate of 300 nl/min: a) for the 6-plex; 2% B for 5 min; 2 to 20 % B in 200 min; 20 to 32% B in 40 min; 32 to 95% B in 1 min; 16 min at 95% B; 95 to 2% B in 1 min; and final wash at 2% B for 20 min. b) for the 16-plex: 2 to 6% B in 5 min; 6 to 25% B in 195 min; 25 to 30% B in 40 min;40 to 95% B in 1 min; 95% B fo 5 min; 95% to 2 % B in 1 min; and a final wash at 2% B for 14 min

All spectra were acquired using an Orbitrap Fusion Lumos mass spectrometer (ThermoScientific), controlled by Xcalibur 2.0 software (ThermoFisher Scientific) and operated in data-dependent acquisition mode. For the 6-plex experiment, survey scans were obtained on the orbitrap analyzer at 120K resolution in the m/z scan range 375 to 1500, with 50 ms maximum injection time and AGC target of 40K. This was followed by tandem mass scan in cycle time dependent mode using an intensity threshold of 50000for ions with charge states 2 to 7, and monisotopic precursor selection, using HCD activation at 35% and detection on the orbitrap analyzer at a resolution of 30K with 360 ms maximum injection time and AGC target of 50K. For the-16 plex experiment, a SPS-MS3 workflow was used.

The cycle consisted of one parent ion scan on the Orbitrap at 60,000 resolution, followed by MSMS scans on the Ion Trap mass analyzer, top speed using CID at 35% energy. A dynamic exclusion of one repeat scans of the same ion with and r-60-s exclusion duration. MS3 scans were performed by synchronous precursor selection (SPS) with ion detection on the orbitrap analyzer at 50K resolution in the m/z scan range 100-500. Peptide identification was done on the MS2 scan and quantification was done on the MS3 scan.

#### Protein identification and quantification

For data analysis all MSMS scans were searched using Protein Discoverer v.2.4 running Sequest HT and a mouse database downloaded from the NCBI data repository (Mus musculus RefSeq database as on 4 November 2019). Full trypsin specificity was defined with a maximum of three missed cleavages and a precursor mass tolerance of 20 ppm and 0.02 Da for fragment tolerance. TMT or MTpro tags for peptide amino terminus and Lys side chain (for 6plex and 16plex, respectively) and carboxymethylation of Cys were used as fixed modifications. Met oxidation; deamidation of Asn and Gln; were used as variable modifications. Phosphorylation of Tyr, Thr or Ser was also selected as a variable modification for phosphopeptide identification. A maximum of three equal modifications and to 4 dynamic modifications was defined. Validation was based on the q-value obtained from Percolator.

### Metabolomic experimental preparation

#### Metabolite extraction

Isolated liver (50 mg) was homogenized in precipitattion solution (8:1:1 Acetonitrile: Methanol: Acetone), vortexed, and kept on ice for 30 minutes. Samples were then centrifuged at 15,000 rcf for 10 minutes to pellet proteins and lipids, supernatant was removed, and dried in Speed Vacuum Centrifuge. Dried metabolites were reconstituted with 50ml 0.1% formic acid and vortexed, placed on ice for 15 minutes, and centrifuged again to remove any residual protein or lipid^10^.

#### Liquid chromatography tandem mass spectrometry

An Agilent HPLC 1100 series was used with a Waters Acquity CSHTM Phenyl-Hexyl 1.7 mM 2.1 _ 50-mm column. A Sciex API 4000 triple quadrupole mass spectrometer with an ESI source was used in positive mode with first scan event a full mass spectrometry scan at 55.0 to 1000 m/z (Boston University Chemical Instrumentation center). For positive polarity, a gradient of 98% buffer A (H2O and 0.1% formic acid) was set at 0.00 to 1.00 minute, buffer B (methanol) increased to 15% at 4.00 minutes, then buffer B increased to

95% to 7.00 minutes then decreased to 2% buffer B at 8.50 minutes through a 10.0 minutes run time. For negative polarity detection of metabolites, a gradient of 98% buffer A and 2% buffer B at 0 to 0.5 minutes, increased buffer B to 95% at 5 minutes, increased buffer B to 98% at 8.5 minutes, and then decreased buffer B to 2% from 9.0 to 10.0 minutes at a flow rate of 0.15 mL/min. In negative polarity mode for analysis, similar parameters were set at 55.0 to 1000 m/z^10^.

### Differential Expression Analysis

#### Transcriptomic

For the RNA-seq data, read counts for each gene were filtered using the ‘filterByExpr’ function, and scaling factors for library sizes were computed with the ‘calcNormFactors’ function using the trimmed mean of M-values (TMM) method^51^, both functions coming from the *edgeR* package ^52-54^ in the R programming language^55^. Counts were transformed to log2-counts per million and observation-level weights were created with limma-voom^56^. Linear models were fit and tests of differential expression were performed using the ‘lmFit’ and ‘eBayes’ functions from the *limma* package^57^.

#### Proteomic & Phospho-proteomic

The proteomic data comes from three datasets. The first contains protein and phosphoprotein abundances for OGT knockout (KO) at 1-2 weeks and was used to compare OGT-KO groups across time points. The second and third datasets, corresponding to protein and phosphoprotein abundances, respectively, contain OGT-KO, OGA-KO, and TMG all at 2 weeks and was used for comparisons between treatments and for the assessment of differential phosphorylation. For all datasets, technical replicates, if present, were aggregated using geometric means. Features with missing data were removed, in lieu of imputation, since proteins with missing values tended to be missing across entire experiments. Dataset 1 contained several independent experiments, which necessitated batch effect correction using the *Combat* method^58^. Each dataset was then log2 transformed and quantile normalized. For all datasets, limma-trend^56^ was used for differential expression analysis with the ‘lmFit’ function and the ‘eBayes’ function with trend=TRUE. Welch’s *t*-test statistics were also computed between log fold changes of common proteins from the proteomic and phospho-proteomic data to determine changes in phosphorylation unrelated to changes in overall abundance of a protein.

#### Metabolomic

The following description was applied to the positive and negative ion modes independently. First, common metabolites were aggregated by the median. Metabolites were filtered based on proportion of point mass values (PMVs) and variability (measured by interquartile range). Metabolites with > 25% of PMVs across samples or those with an IQR of 0 were removed from the analysis (n=285). Values were log2 transformed and quantile normalized. MetaboAnalyst^59^ was used to map names to PubChem IDs^60^, and metabolites that did not map were removed (n=12). The limma-trend method^56^ was used for differential expression analysis by using the ‘lmFit’ function and the ‘eBayes’ function with trend=TRUE.

#### Equivalent Change Index

The Equivalent Change Index (ECI) measures the degree to which the change in expression for a gene is similar or different between two experiments^61^. It involves taking the ratio of log fold changes coming from two treatment-control comparisons. The ECI allows us to overcome confounding due to batch effects when attempting to compare the effects of two treatments coming from different experiments. Several types of ECIs were calculated for each “Omic” type. In what follows, *T* denotes transcriptomic, *P* denotes proteomic, *Ph* denotes phospho-proteomic, and *M* denotes metabolomic. ECIs were calculated between 1-week OGT-KO and 2-week OGT-KO (for *T*, *P*, *Ph*, and *M*), 1-week OGA-KO and 2-week OGA-KO (for *T* and *M*), and between 1-week TMG and 2-week TMG (for *T*). ECIs were also calculated between all pairwise combinations of OGT-KO, OGA-KO and TMG, all at 2 weeks (for *T, P, Ph,* and *M*). For the metabolomic ECI calculations, the log fold changes of metabolites present in both positive and negative mode were first aggregated by the median.

#### Molecular Interaction Networks

The *Mus musculus* protein-protein interaction (PPI) network was downloaded from STRING v.11.5^62^, which included both physical and functional PPIs. Only edges with a combined score of 0.7 or greater were kept to reduce potential false-positive interactions. Gene symbols were mapped to Ensembl peptide IDs using the *biomaRt* R package^63^. Only features that were present in at least one experiment and that were successfully mapped to an Ensembl peptide ID were kept. After filtering and taking the largest connected component, the PPI network consisted of 11,318 nodes and 169,451 edges.

The metabolite-metabolite interaction (MMI) network was downloaded from STITCH v.5.0^64^. We applied similar pre-processing steps taken for the PPI network. Only edges with an edge score greater than or equal to 0.7 were kept. Metabolites whose PubChem IDs didn’t map to any experimental data were removed. This resulted in a connected network of 187 nodes and 1,683 edges.

#### Active Module Identification

Active Module Identification (AMI) is the search for subnetworks whose nodes are ‘active’ under, or biologically relevant to, the experimental conditions of interest, *e.g.*, genes that show large changes in expression between treatment and control samples^65^, The AMI method used in this study is AMEND (Active Module identification with Experimental data and Network Diffusion), a previously introduced algorithm that searches for subnetworks of features with large experimental values and high connectivity^6^. For the algorithm settings, the final module size was set to 50 for transcriptomic, proteomic, and phospho-proteomic data, and to 20 or 25 for the metabolomic data. Adjacency matrices were normalized by node degree. Complete edge lists for all AMEND modules are provided in Extended Table 2A-OO.

#### Degree Bias

Due to various technical and study biases associated with PPI ascertainment, some proteins may have an inflated or deflated degree relative to others in the PPI network^66^. This is problematic since many network methods, including RWR, learn from node degree. In the context of RWR, degree influence for a node manifests itself in the stationary distribution of the transition matrix, whose probabilities are purely a function of degree when the restart parameter equals zero. Mitigating degree influence translates into squeezing these probabilities towards a global mean, the success of which can be measured by the entropy of the stationary distribution, with larger entropy meaning a more homogeneous stationary distribution. To mitigate degree bias, we utilize the inflation operator from Markov Clustering^67^ where values in each row are raised to a certain power greater than 1, with the exponent in each row being a positive linear function of the stationary distribution probability of the node associated with that row. This linear function has an intercept of 1 and a slope chosen such that the entropy of the stationary distribution of the resulting transition matrix is maximized (using a grid search). The inflation operator causes incoming edge weights of a node to be displaced to the edges of its neighbors in proportion to the amount of degree influence on that node. Since the row-wise inflation operator will perturb column sums away from one, all columns are re-normalized to sum to one after inflation to ensure left-stochasticity of the transition matrix. This inflation-normalization procedure effectively decreases the influence of degree on diffusion scores, represented by a decrease in correlation between diffusion scores and node degree (**Extended Table 3**).

#### Visualization

Subnetworks were visualized using the Cytoscape software^68, 69^. To connect subnetworks to functional enrichment results, we performed within-module cluster annotation. First, the Louvain topological clustering algorithm was applied to a module to get clusters^70^. For each cluster, we chose a representative pathway by counting the number of nodes in the cluster that each pathway contains and picking the pathway with the largest count. Clusters with the same representative pathway were merged.

#### Over-representation Analysis

Over-representation Analysis (ORA) was used to functionally characterize the subnetworks from AMEND. For the transcriptomic, proteomic, and phospho-proteomic subnetworks, pathways from the Reactome database were used^71^, with the FDR controlled at 0.05 following the Benjamini-Hochberg procedure for multiple testing^71^. Metabolic pathways from KEGG^72^ were used for the metabolomic subnetworks. Given the exploratory nature of the bioinformatical analysis and the weaker signal of the metabolomic data, we opted for no adjustment for multiple testing while controlling the false-positive rate at 0.05 for the metabolomic ORA analysis. ORA results are displayed here as directed, acyclic graphs, wherein a pathway is represented by a node with a directed edge going to the larger pathway within which it is (partially) nested. To achieve this, a similarity matrix is computed containing all pairwise measures of nested-ness between significant pathways from ORA. The extent to which a pathway with feature set A is nested within a pathway with feature set B is given by f(A,B)=(|A∩B|)/(min(|A|,|B|)). After applying a threshold to these values, the similarity matrix is treated as an adjacency matrix from which a directed graph is created by only keeping edges going from a smaller pathway to a larger one. For interpretability, only the longest simple directed path between each pair of vertices is kept.

#### Gene Set Enrichment Analysis

We used Gene Set Enrichment Analysis (GSEA) to identify pathways associated with changes in expression between treatment and control groups^73^. GSEA was applied to all sets of log fold changes from differential expression analysis. Both GSEA and ORA were implemented in R using the *fgsea* package (2023, fgsea: Fast Gene Set Enrichment Analysis, Github).

### Flow cytometry

PH was performed on OGT floxed, OGT-KO, OGA-floxed, and OGA-KO mice, and 14 days after PH hepatocytes were isolated by Cell Isolation Core at KUMC. Hepatocytes were permeabilized in cold 70% ethanol overnight at 4^0^C. The next day, cells were stained with propidium iodide (Thermo Fisher Scientific F10797) for 30 minutes at room temperature in the dark. Ploidy analysis was performed using the LSR II flow cytometer (BD Biosciences) and FACSDiva software by the Flow Cytometry Core Facility at KUMC.

## Acknowledgements

C.S discloses support of this work from NIH R01AAG06422. Support for this work from NIH R01 DK0198414 was provided to U.A. Support was also provided through the University of Kansas Alzheimer’s Disease Research Center P30AG072973.

---

**Extended Figure 1:**
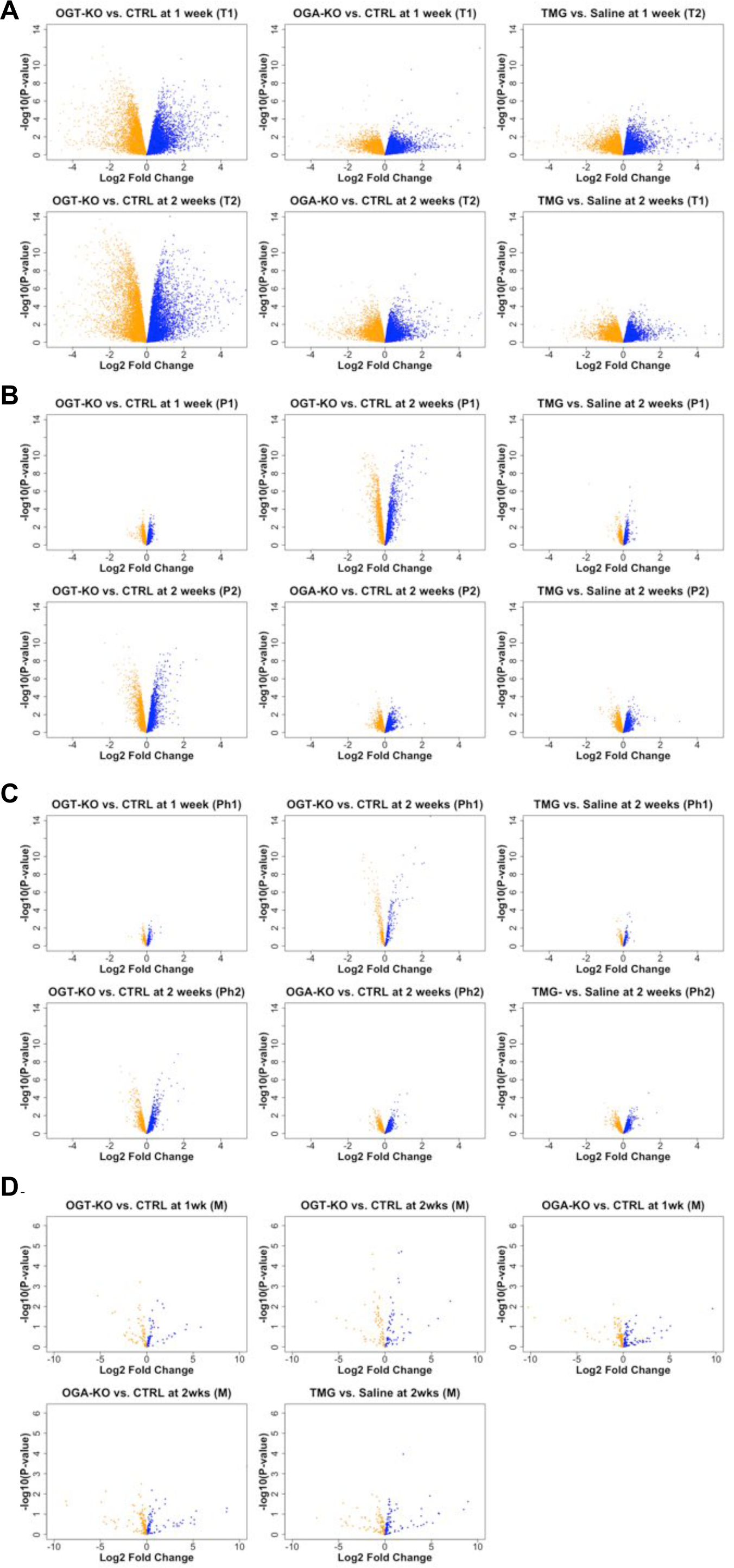
Differential expression of transcripts, proteins, and metabolites after multi-omics analysis: Volcano plots were generated in R by plotting log fold changes against -log10-transformed unadjusted p-values from differential expression analyses. **A:** Volcano plots of transcriptomic changes in OGT KO after 1 and 2 weeks, OGA KO after 1 and 2 weeks, and TMG treatment after 1 and 2 weeks. **B**: Volcano plots of proteomic changes in OGT KO after 1 and 2 weeks (experimental group 1 and experimental group 2), OGA KO after 2 weeks (experimental group 1 and experimental group 2), and TMG treatment after 2 weeks (experimental group 1 and experimental group 2). **C**: Volcano plots of phospho-proteomic changes in OGT KO after 1 and 2 weeks (experimental group 1 and experimental group 2), OGA KO after 2 weeks (experimental group 1 and experimental group 2), and TMG treatment after 2 weeks (experimental group 1 and experimental group 2). **D.** Volcano plots of metabolite changes in OGT KO after 1 and 2 weeks, OGA KO after 1 and 2 weeks, and TMG treatment after 2 weeks.

**Extended Figure 2:**
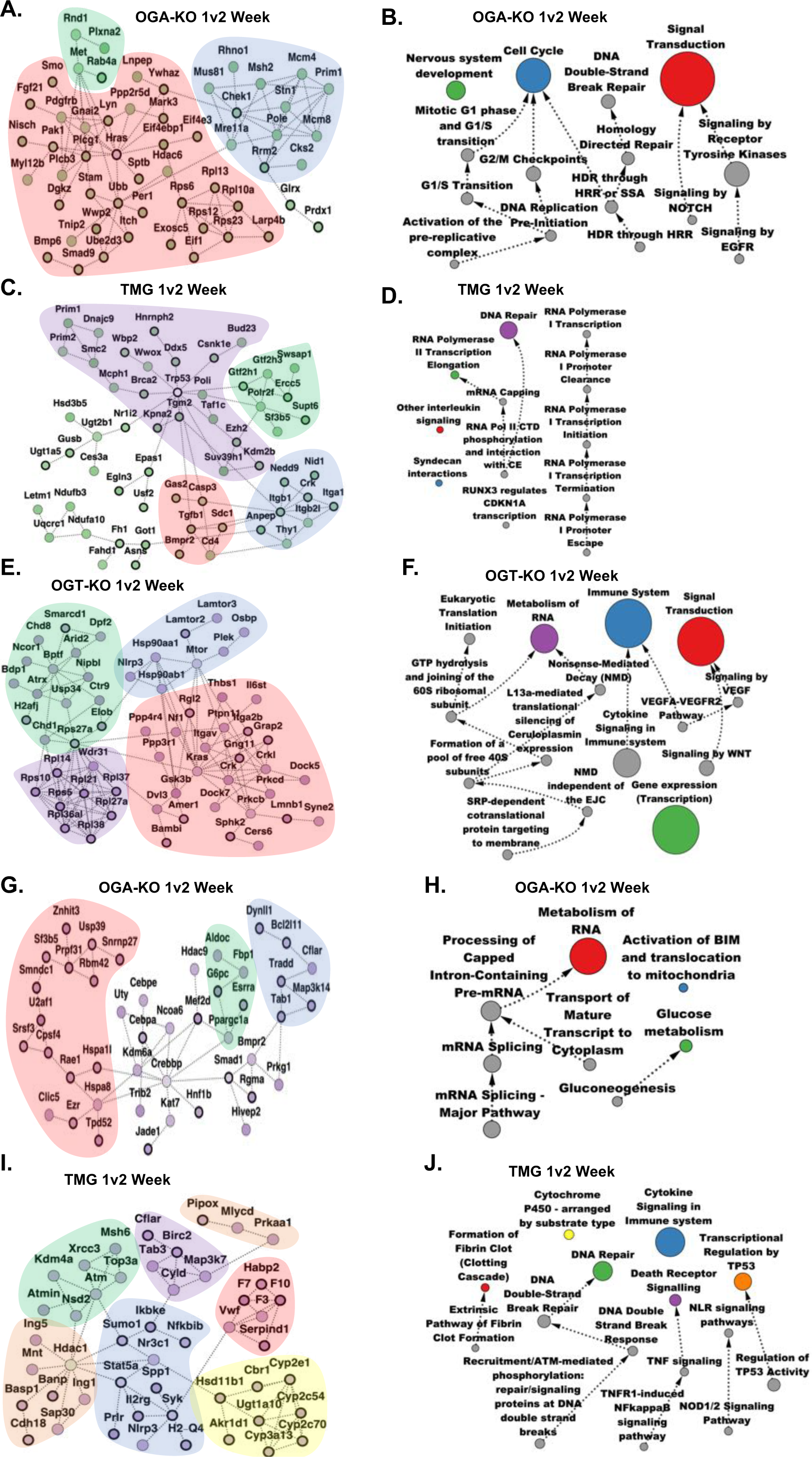
Transcriptomic subnetworks from AMEND, along with pathways from Overrepresentation Analysis (ORA) represented as a directed acyclic graph with longitudinal sampling: Each AMEND subnetwork compares log fold changes between two treatment-control comparisons using the Equivalent Change Index (ECI). Green nodes represent equivalent change between treatment groups 1 and 2, with a bold border signifying that both treatment groups were up-regulated compared to their controls and a thin border signifying that both treatment groups were down-regulated compared to their controls. Purple nodes represent inverse change between treatment groups 1 and 2, with a bold border signifying that treatment 1 was up-regulated while treatment 2 was down-regulated, and vice-versa for a thin border. “T” = transcriptomic,”+” = AMEND selects for positive ECI nodes (equivalent change). Clusters within each subnetwork are colored corresponding to the pathway associated with the nodes in that cluster. Treatment 1 and treatment 2 correspond to the first and second treatment listed in the panel description, respectively. Arrows in the ORA networks show the direction of nestedness (i.e., the source node is nested within the target node) within the directed acyclic graph and the sizes of the pathways are represented by the relative sizes of the nodes. The color of the node reflects the AMEND protein interaction network. Only the top 15 pathways, ranked by adjusted p-value, are shown. **A**: AMEND module, OGA-KO 1 week vs. 2 week (T)(+), **B**: ORA pathways, OGA-KO 1 week vs. 2 week (T)(+), **C**: AMEND module, TMG 1 week vs. 2 week (T)(+), **D**: ORA pathways, TMG 1 week vs. 2 week (T)(+), **E**: AMEND module, OGT-KO 1 week vs. 2 week (T)(-), **F**:ORA pathways, OGT-KO 1 week vs. 2 week (T)(-), **G**: AMEND module, OGA-KO 1 week vs. 2 week (T)(-), **H**: ORA pathways, OGA-KO 1 week vs. 2 week (T)(-), **I**: AMEND module, TMG 1 week vs. 2 week (T)(-), **J**: ORA pathways, TMG 1 week vs. 2 week (T)(-).

**Extended Figure 3:**
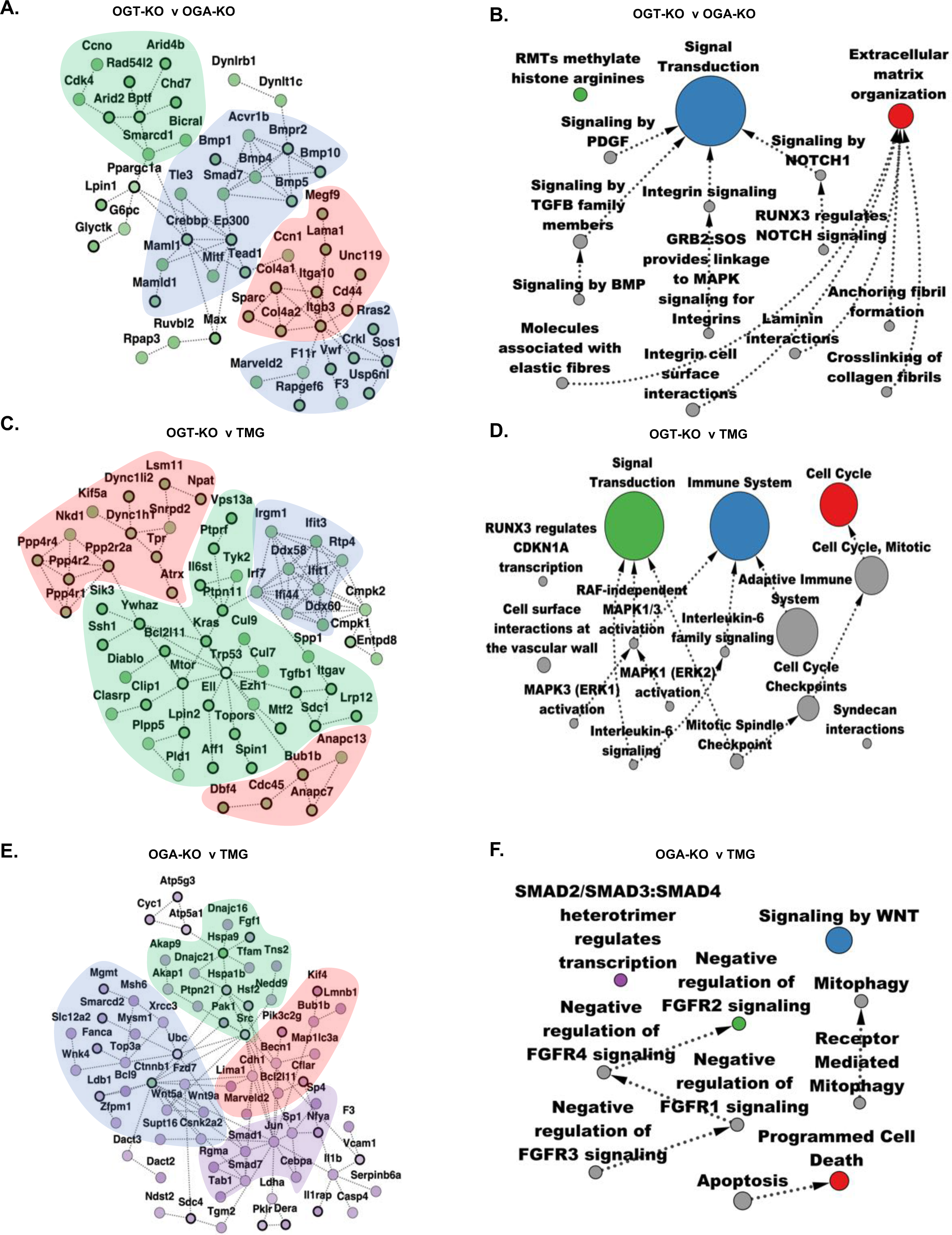
Transcriptomic subnetworks from AMEND, along with pathways from Overrepresentation Analysis (ORA) represented as a directed acyclic graph with heterogeneous sampling: “T” = transcriptomic, “+” = AMEND selects for positive ECI nodes (equivalent change), “-” = AMEND selects for negative ECI nodes (inverse change). Clusters within each subnetwork are colored corresponding to the pathway associated with the nodes in that cluster. Treatment 1 and treatment 2 correspond to the first and second treatment listed in the panel description, respectively. Arrows in the ORA networks show the direction of nestedness (i.e., the source node is nested within the target node) within the directed acyclic graph within the directed acyclic graph and the sizes of the pathways are represented by the relative sizes of the nodes. The color of the node reflects the AMEND protein interaction network. Only the top 15 pathways, ranked by adjusted p-value, are shown. **A**: AMEND module, OGT-KO vs. OGA-KO at 2 weeks (T)(+), **B**: ORA pathways, OGT-KO vs. OGA-KO at 2 weeks (T)(+), **C**: AMEND module, OGT-KO vs. TMG at 2 weeks (T)(+), **D**: ORA pathways, OGT-KO vs. TMG at 2 weeks (T)(+), **E**: AMEND module, OGA-KO vs. TMG at 2 weeks (T)(-), **F**: ORA pathways, OGA-KO vs. TMG at 2 weeks (T)(-).

**Extended Figure 4:**
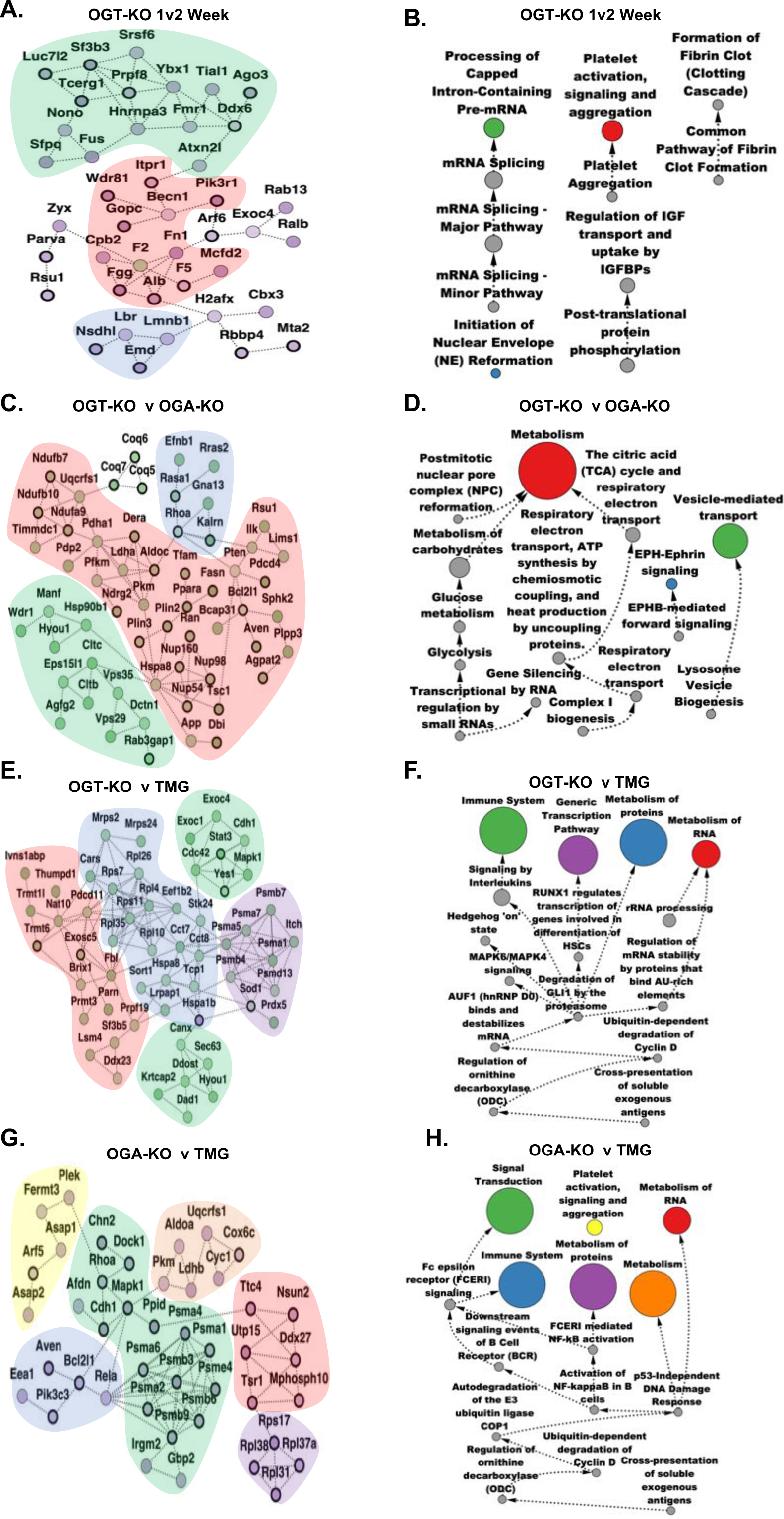
Proteomic subnetworks from AMEND, along with pathways from Overrepresentation Analysis (ORA) represented as a directed acyclic graph with longitudinal or heterogeneous sampling: “P” = proteomic,”+” = AMEND selects for positive ECI nodes (equivalent change) “-” = AMEND selects for negative ECI nodes (inverse change). Clusters within each subnetwork are colored corresponding to the pathway associated with the nodes in that cluster. Treatment 1 and treatment 2 correspond to the first and second treatment listed in the panel description, respectively. Arrows in the ORA networks show the direction of nestedness (i.e., the source node is nested within the target node) within the directed acyclic graph and the sizes of the pathways are represented by the relative sizes of the nodes. The color of the node reflects the AMEND protein interaction network. Only the top 15 pathways, ranked by adjusted p-value, are shown. **A**: AMEND module, OGT-KO 1 week vs. 2 week (P)(-), **B**: ORA pathways, OGT-KO 1 week vs. 2 week (P)(-), **C**: AMEND module, OGT-KO vs. OGA-KO at 2 weeks (P)(+), **D**: ORA pathways, OGT-KO vs. OGA-KO at 2 weeks (P)(+), **E**: AMEND module, OGT-KO vs. TMG at 2 weeks (P)(+), **F**: ORA pathways, OGT-KO vs. TMG at 2 weeks (P)(+), **G**: AMEND module, OGA-KO vs. TMG at 2 weeks (P)(-), **H**: ORA pathways, OGA-KO vs. TMG at 2 weeks (P)(-).

**Extended Figure 5:**
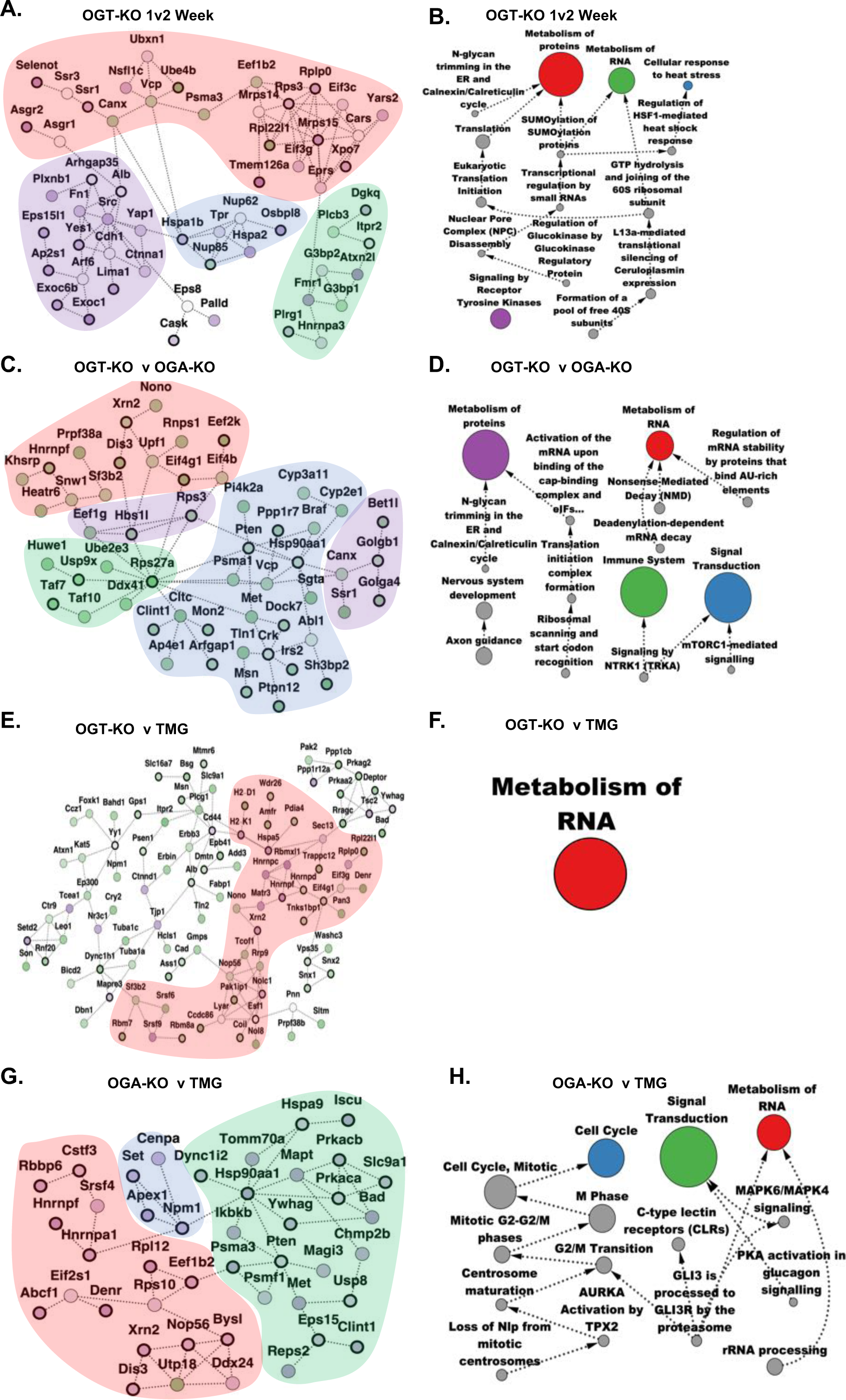
Phospho-proteomic subnetworks from AMEND, along with pathways from Overrepresentation Analysis (ORA) as a directed acyclic graph with longitudinal or heterogeneous sampling: “Ph” = phospho-proteomic,”+” = AMEND selects for positive ECI nodes (equivalent change) “-” = AMEND selects for negative ECI nodes (inverse change). Clusters within each subnetwork are colored corresponding to the pathway associated with the nodes in that cluster. Treatment 1 and treatment 2 correspond to the first and second treatment listed in the panel description, respectively. Arrows in the ORA networks show the direction of nestedness (i.e., the source node is nested within the target node) within the directed acyclic graph and the sizes of the pathways are represented by the relative sizes of the nodes. The color of the node reflects the AMEND protein interaction network. Only the top 15 pathways, ranked by adjusted p-value, are shown. **A**: AMEND module, OGT-KO 1 week vs. 2 week (Ph)(-), **B**: ORA pathways, OGT-KO 1 week vs. 2 week (Ph)(-), **C**: AMEND module, OGT-KO vs. OGA-KO at 2 weeks (Ph)(+), **D**: ORA pathways, OGT-KO vs. OGA-KO at 2 weeks (Ph)(+), **E**: AMEND module, OGT-KO vs. TMG at 2 weeks (Ph)(+), **F**: ORA pathways, OGT-KO vs. TMG at 2 weeks (Ph)(+), **G**: AMEND module, OGA-KO vs. TMG at 2 weeks (Ph)(-), **H**: ORA pathways, OGA-KO vs. TMG at 2 weeks (Ph)(-).

**Extended Figure 6:**
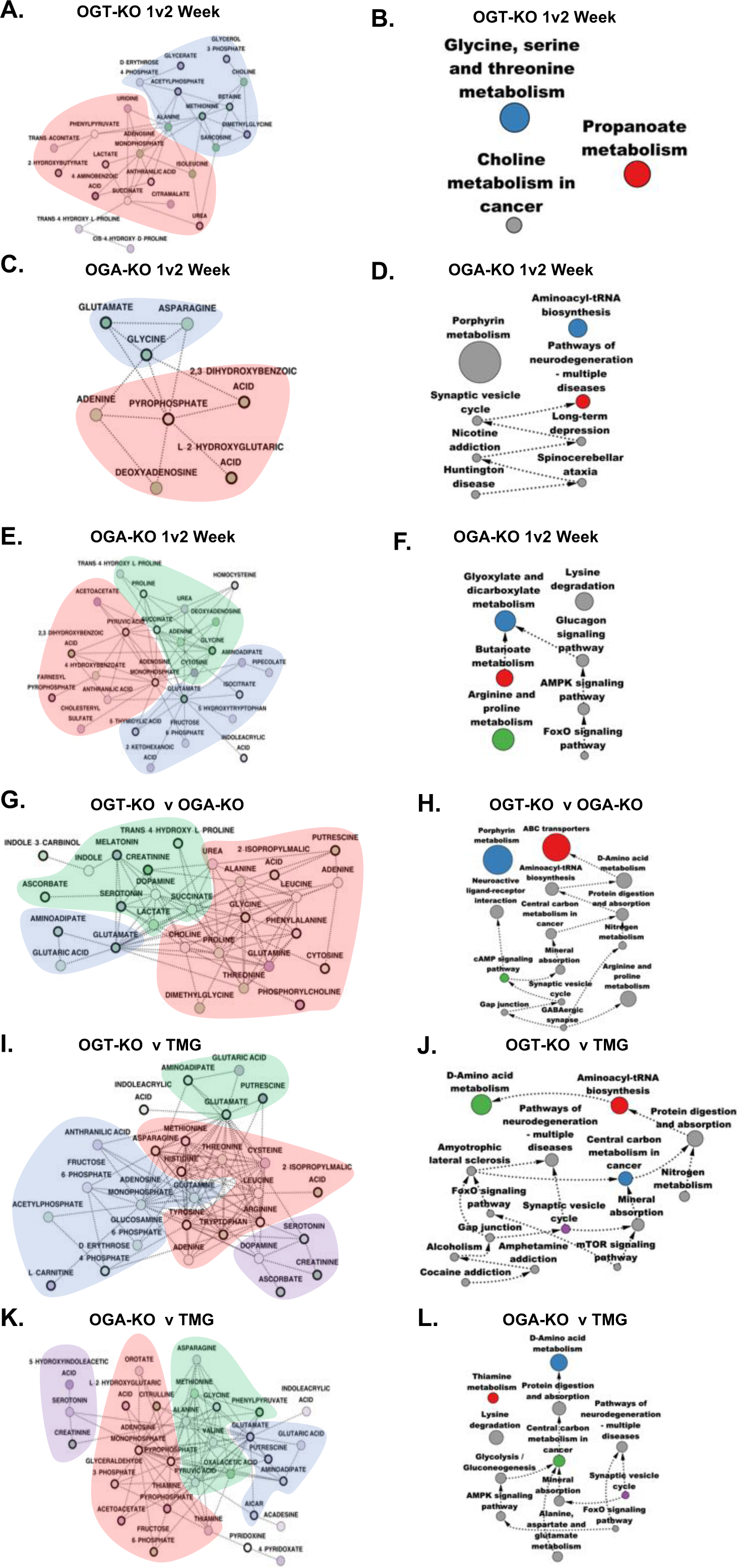
Identification of metabolomic subnetworks from using AMEND and Overrepresentation Analysis (ORA) from longitudinal or heterogeneous sampling: “M” = metabolomic, “+” = AMEND selects for positive ECI nodes (equivalent change), “-” = AMEND selects for negative ECI nodes (inverse change). Clusters within each subnetwork are colored corresponding to the pathway associated with the nodes in that cluster. Treatment 1 and treatment 2 correspond to the first and second treatment listed in the panel description, respectively. Arrows in the ORA networks show the direction of nestedness (i.e., the source node is nested within the target node) and the sizes of the pathways are represented by the relative sizes of the nodes. Only the top 15 pathways, ranked by p-value, are shown**. A**: AMEND module, OGT-KO 1 week vs. 2 week (M)(-), **B**: ORA pathways, OGT-KO 1 week vs. 2 week (M)(-), **C**: AMEND module, OGA-KO 1 week vs. 2 week (M)(+), **D**: ORA pathways, OGA-KO 1 week vs. 2 week (M)(+), **E**: AMEND module, OGA-KO 1 week vs. 2 week (M)(-), **F**: ORA pathways, OGA-KO 1 week vs. 2 week (M)(-) **G**: AMEND module, OGT-KO vs. OGA-KO at 2 weeks (M)(+), **H**: ORA pathways, OGT-KO vs. OGA-KO at 2 weeks (M)(+), **I**: AMEND module, OGT-KO vs. TMG at 2 weeks (M)(+), **J**: ORA pathways, OGT-KO vs. TMG at 2 weeks (M)(+), **K**: AMEND module, OGA-KO vs. TMG at 2 weeks (M)(-), **L**: ORA pathways, OGA-KO vs. TMG at 2 weeks (M)(-).

**Extended Figure 7:**
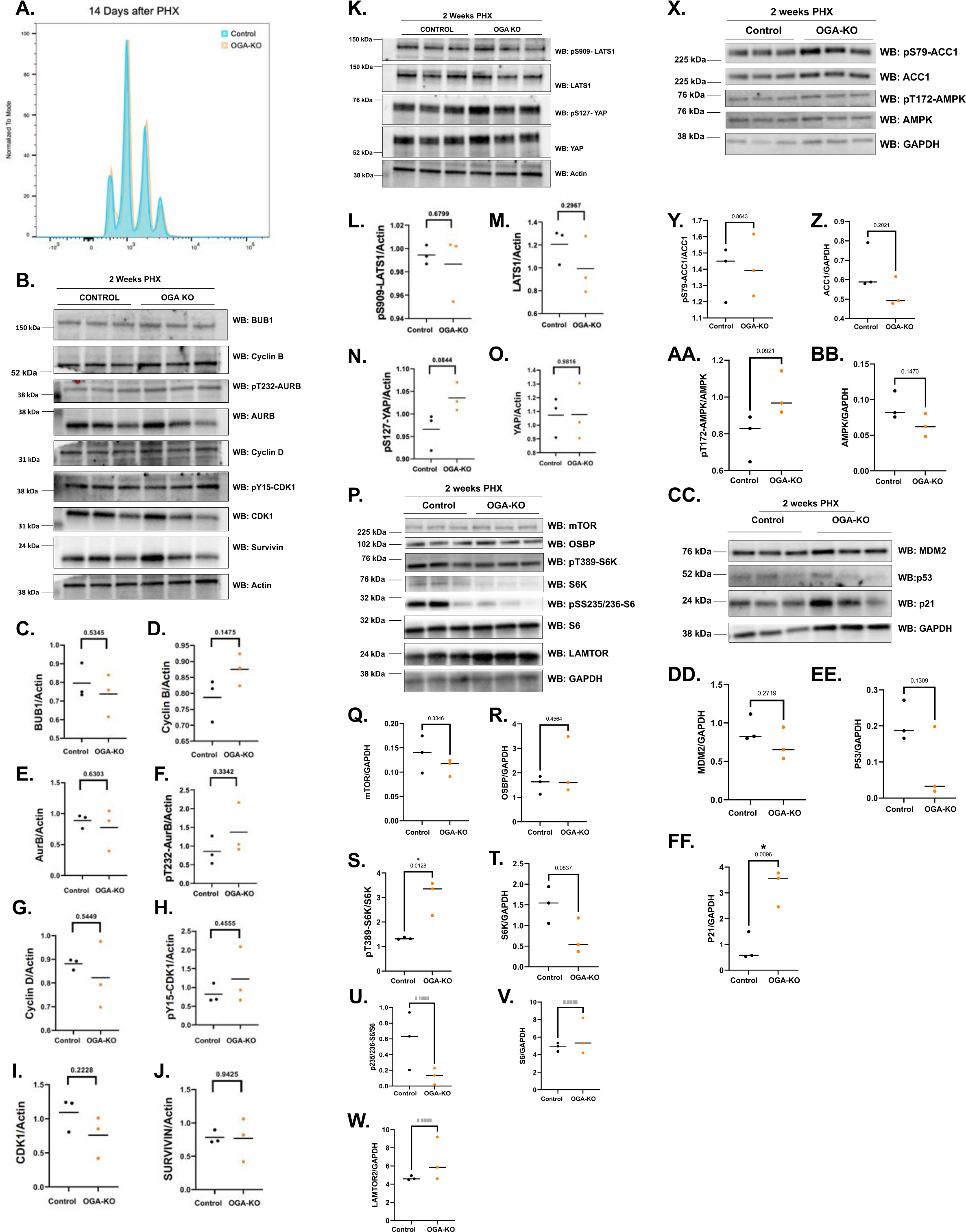
OGA knockout livers do not present with aneuploidy after PHX: A: Flow cytometry of OGA KO livers show no changes in DNA content after PHX (blue is no PHX, red is after PHX). **B:** Western blot analysis on mitotic proteins. Actin is used as a load control (n=3). **C-J:** Densitometry of western blot samples. * = p value less than 0.05. **K:** Western blot of HIPPO signaling pathway proteins (n=3). Actin is used as a load control **L-O:** Densitometry of western blot samples. * = p value less than 0.05. **P:** Western blot of mTOR signaling pathway proteins (n=3). GAPDH is used as a load control (n=3). **Q-W:** Densitometry of western blot samples. * = p value less than 0.05. **X:** Western blot of AMPK signaling pathway proteins (n=3). GAPDH is used as a load control (n=3). **Y-BB:** Densitometry of western blot samples. * = p value less than 0.05. **CC:** Western blot of p53 signaling pathway proteins (n=3). GAPDH is used as a load control (n=3). **DD-FF:** Densitometry of western blot samples. * = p value less than 0.05.

## Notes

### Competing Interest Statement

The authors have declared no competing interest.

